# TOPAS: phosphoproteome data analysis and decision support platform for molecular tumor boards

**DOI:** 10.64898/2026.07.08.737143

**Authors:** Cecilia Bang Jensen, Amirhossein Sakhteman, Firas Hamood, Annika Schneider, Julia Woortman, Maria-Veronica Teleanu, Peter Horak, Florian P. Bayer, Christoph Stange, Jennifer Huellein, Daniel Hübschmann, Anton G. Henssen, Stefan Fröhling, Bernhard Kuster, Matthew The

**Author notes:** these authors contributed equally (Cecilia Bang Jensen, Amirhossein Sakhteman, Firas Hamood, Annika Schneider).

## Abstract

The molecular tumor board (MTB) is central to precision oncology, providing personalized treatment recommendations based on molecular profiles of patient tumors. Genomics is instrumental for MTBs but often fails to identify clinically actionable targets, a gap that phosphoproteomics can fill. We present the tumor proteome activity status (TOPAS) platform, an end-to-end analysis pipeline that converts terabytes of phosphoproteomic data into patient-specific reports for MTB discussions, focusing on clinically relevant signaling linked to oncogenic mechanisms and therapeutic targets. Designed to scale with growing cohorts, the platform integrates data from 1,998 tumor samples to support patient- and cohort-level hypothesis generation. A web portal handles quality control, calculates TOPAS scores, identifies tumor antigens and immune checkpoints, and offers interactive analyses of differential protein abundance and outlier detection. The TOPAS platform is open source, addresses a critical unmet need and facilitates broader adoption of phosphoproteomics in precision oncology in the future.

---

Retrospective molecular tumor profiling initiatives such as The Cancer Genome Atlas (TCGA)^1^ and the Clinical Proteomic Tumor Analysis Consortium (CPTAC)^2^ have demonstrated conceptual benefits of analyzing pan-cancer cohorts over single cancer types. Examples include the identification of molecular signatures, cancer (immune) subtypes, or signaling patterns that span entities or demarcate certain entities^3, 4^. The pan-cancer perspective is particularly relevant for the analysis of patients suffering from the large variety and molecularly heterogeneous group of rare or otherwise hard-to-treat cancers. For many patients, standard-of-care systemic therapy options are either limited or not available at all^5, 6^. Molecular tumor boards (MTBs) aim to make therapeutic recommendations for such cases based on their individual genomic and, sometimes, transcriptomic profiles^5, 7^. Despite noteworthy successes, identifying or predicting actionable oncogenic drivers from nucleotide data remains challenging, and clinical trials conducted so far only showed moderate patient benefit^5, 6, 8, 9^. Because many cancers are driven by aberrant protein expression or signaling pathway activity, proteomics and phosphoproteomics (from here collectively referred to as phosphoproteomics) are important additional data layers. In an accompanying paper^10^, we demonstrate that adding this component to MTBs is not only technically feasible but also enhances treatment recommendations across many tumor entities. Here, we present the enabling end-to-end and open-source data analysis platform that turns raw phosphoproteomic data into patient-specific reports to support decision-making in MTBs and enables gaining novel insights into the tumor biology of rare cancers such as alveolar rhabdomyosarcoma (ARMS) in a robust, rapid, and scalable manner.

## From raw phosphoproteome data to MTB reports

Integrating phosphoproteome data into prospective weekly MTB meetings required the development of a new and bespoke data processing pipeline that followed three major design principles (**Fig. 1a**).

**Fig. 1:**
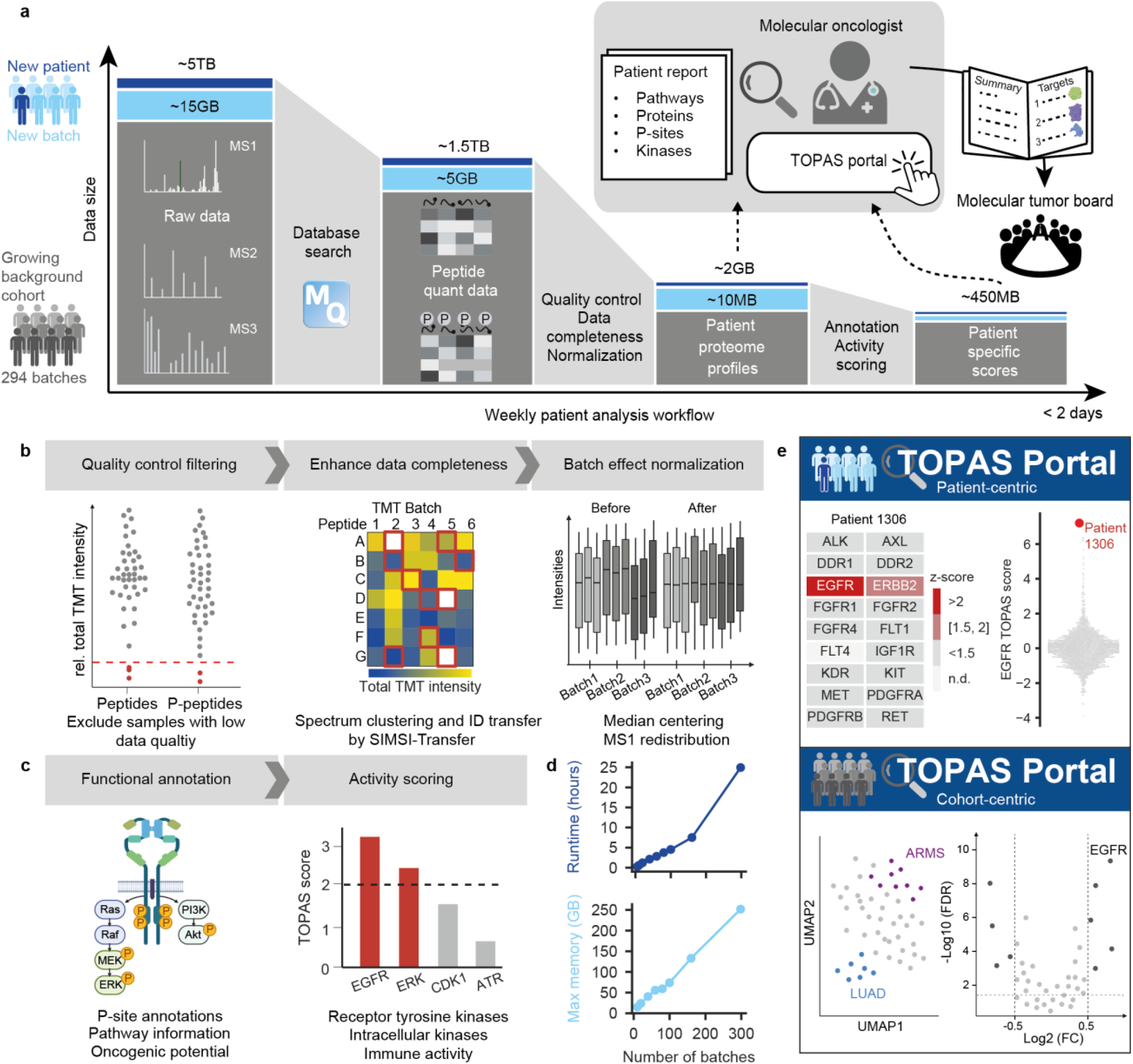
End-to-end phosphoproteome data analysis platform. (a) Schematic of how the data analysis pipeline condenses raw mass spectrometric protein and phosphopeptide (p-peptide) identification and quantification data into patient-specific reports and scores of tumor activity, and imports these into an interactive web-based data portal where molecular oncologists analyze the results to produce a summary for MTB discussions (MQ, MaxQuant software, TOPAS, tumor proteome activity status). (b) Along the path depicted in (a), the output of MaxQuant is assessed by quality metrics (red line, minimal threshold for acceptance), enhanced for data completeness (TMT, tandem mass tags, ID, peptide identification), and reduced for batch effects (MS1, mass spectrometric peptide intensity). (c) Patient phosphoproteome profiles are annotated using prior knowledge from databases and the scientific literature, and scored for signaling activity (black dotted line indicates significance threshold). (d) Runtime and memory usage with increasing number of batches. (e) Processed data is imported into a web-based data portal where it can be interrogated in a patient- and cohort-centric manner.

First, it needed to place data collected for an individual patient into an ever-growing background cohort of samples from the same or different tumor entities. This design increases statistical power and confidence in the biological and clinical interpretation of the profiles over time. The current data set comprises ∼4 TB of raw mass spectrometry (MS) data (17,640 MS runs) collected from 1,998 tumor samples from 1,852 patients. Analyzing all data for peptide and protein identification and quantification in a single step each week was not feasible with reasonable computing resources. Instead, each batch of 8 or 9 patients plus two or three common cell line quality control (QC) samples (multiplexed by stable isotope labeling using tandem mass tags, TMT, **Methods**) was searched separately. Results from 294 such batches were combined using mass spectra clustering by SIMSI-Transfer to reduce missing data between batches^11^ and applying the picked protein group approach^12^ to control the false discovery rate (FDR) of protein identification at 1%. This led to the quantification of 13,069 protein-coding genes and 148,224 phosphopeptides, of which 8,192 and 22,414 respectively were detected in ≥50% of all patient samples (**Extended Data Fig. 1a**). Samples with poor TMT labeling (<90%), phosphopeptide enrichment (<80%), or a summed TMT intensity of 5-fold below QC samples in the same batch were removed (**Fig. 1b, Extended Data Fig. 1b, Methods**) and normalization was applied to reduce batch effects (**Fig. 1b, Methods**).

Second, to aid molecular oncologists with data interpretation, the data volume per patient had to be reduced to the essentials and analyzed using easily explainable methods rather than potentially more powerful, but less intuitively accessible ones. For instance, whenever available, proteins and phospho-sites of interest were annotated using prior knowledge from databases and the scientific literature. In addition, tumor proteome activity status (TOPAS) scores and kinase substrate phosphorylation scores were calculated for 18 receptor tyrosine kinases (RTKs) and 28 intracellular kinases (ICKs) (**Fig. 1c**). The data processing pipeline takes about one day to create >2,000 annotated patient-specific reports from the raw data using off-the-shelf computing hardware (**Fig. 1d**).

Third, the data needed to be accessible in a simple fashion. This was achieved by importing abundance values, annotations and phosphoproteome scores into an interactive TOPAS Portal (**Fig. 1e**) that molecular oncologists use in tandem with a patient-specific tabular report (<5 MB, downloadable from the portal, **Extended Data Table 1**) to obtain an in-depth appreciation of potentially dysregulated protein expression and pathway signaling in a patient’s tumor. This information is summarized into a single page and discussed alongside the genomics and transcriptomics data for the same patient in the weekly MTB.

The TOPAS platform can also be applied to analyzing cohort data from other sources. Because the patient data of the current study requires controlled access (see data availability), we demonstrate this capability by public data from the CPTAC data portal, comprising 153 breast, 225 lung and 153 Uterine Corpus Endometrial Carcinoma (UCEC) tumor samples and healthy controls^2^. Additionally, we imported all 44 sarcoma samples from a recently published pan-cancer proteome atlas^13^ to demonstrate compatibility with label-free quantification and data independent acquisition (DIA) data. We note that only a subset of the functionality is available for this data source, as the phosphoproteome was not acquired.

## Measuring and managing batch effects

Addressing technical variation is critical when integrating data over the course of months to years. Five MS instruments were deployed on the project, and we observed performance drifts over time, leading to batch effects (**Fig. 2a)** that were also apparent at the level of tumor entities and individual proteins (**Extended Data Fig. 2**). The conventional approach to TMT batch correction is to use a common reference sample (here two or three cell line QC samples) contained in every batch^14^. However, this approach would have resulted in missing >5,000 proteins detected in patients, which were insufficiently abundant in the QC samples, an effect exacerbated by the biological heterogeneity of a pan-cancer cohort. Moreover, batch-correction methods run the risk of introducing artifacts which can complicate data interpretation^15^. Instead, we addressed batch effects as follows (**Methods**): To overcome missing data in QC samples, we computed quasi-label-free quantification (LFQ) intensities^2, 16^ for any sample (QC or patient) by distributing the precursor intensity (produced in the MS1 spectrum) across the relative contributions of the TMT reporter fragment ion intensities (produced in the MS3 spectrum; **Extended Data Fig. 3a**). This approach has the additional benefit of allowing comparison of cohorts that do not share a common reference sample. For protein quantification, we observed that summing all peptide intensities of a protein produced substantial batch effects (**Extended Data Fig. 3b**). This is because peptides identifying a particular protein in different samples are not necessarily the same^17^. Using the MaxLFQ approach^17^ on the quasi-LFQ intensities for protein quantification alleviated these issues. We found that median z-scored MS intensities of all proteins for a patient were correlated with the number of proteins identified in a given batch. This effect was minimized by using only proteins detected in ≥70% of samples for median centering normalization (**Extended Data Fig. 3c**).

**Fig. 2:**
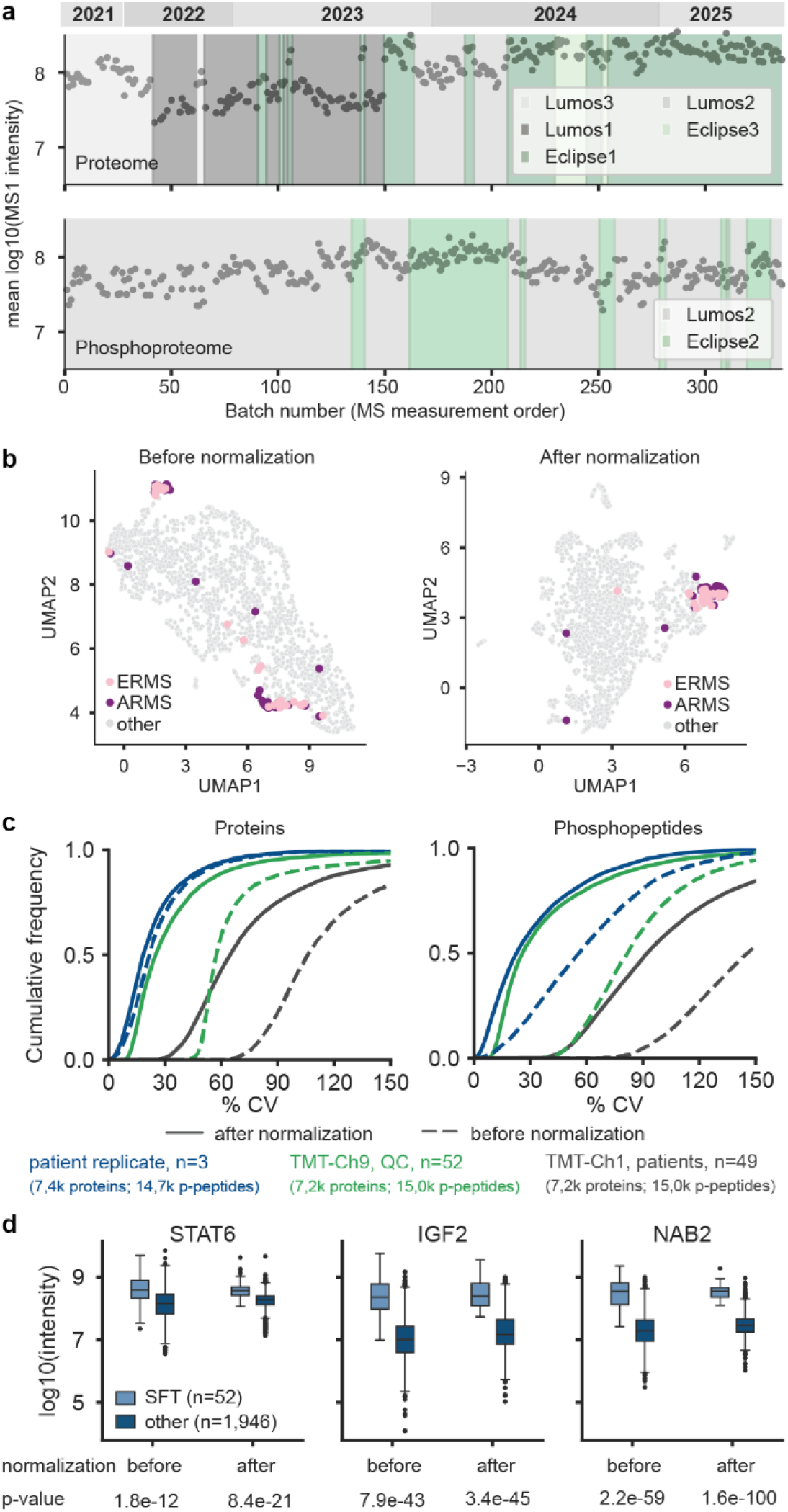
Data normalization across sample batches. (a) MS1 signal intensity drift and differences between mass spectrometers for proteomes (top) and phosphoproteomes (bottom) over time. (b) UMAPs analysis of protein abundance of all patient samples before (left) and after (right) normalization. (c) Cumulative frequency plots showing the effect of data normalization on quantitative precision (coefficient of variation, CV) of technical replicates of identical patient samples (n=3), QC samples (n=52), or diverse patient samples (n=49; processed in different batches) on protein and phosphopeptide level. (d) Box plots showing the abundance of exemplary biomarkers for Solitary fibrous tumors (SFT: n=49 for STAT6, IGF2, NAB2) compared to all other patients (Other: STAT6 n=1,949, IGF2 n=1,928, NAB2 n=1,896) samples before and after normalization.

Combining these three steps led to improved clustering of samples from similar entities such as alveolar rhabdomyosarcoma (ARMS) and embryonal rhabdomyosarcoma (ERMS) ( **Fig. 2b, Extended Data Fig. 2a, b**). Fit-for-purpose repeatability across batches was assessed for full process replicates of the same patient sample (n=3, 18% median CV for proteins, 23% for phosphopeptides; **Fig. 2c**, 7 additional patients in **Extended Data Fig. 4**) as well as QC sample replicates (n=52, 25% median CV for proteins and 26% for phosphopeptides; **Fig. 2c**). Across patients measured in the same TMT channel but across 49 different batches, median CVs were much larger (65% for proteins and 92% for phosphopeptides), demonstrating that technical and biological abundance variations could be distinguished for the majority of cases. These figures of merit provide important guidance for assessing differential abundance data between patients or cohorts. Data normalization also improved the interpretability of biological effects by reducing the within-entity standard error of the mean (median reduction of 45% for proteins and 37% for phosphopeptides, **Methods**). This is exemplified by an increased statistical significance of differential abundance of NAB2, STAT6 and IGF2, known biomarkers for Solitary Fibrous Tumors (SFT)^18^ (**Fig. 2d**) as well as EWSR1, FLI1, CD99, all biomarkers for Ewing Sarcoma (ES)^19^ **(Extended Data Fig. 2c**). Similar benefits were observed for phosphopeptides of interest in ES (**Extended Data Fig. 2d**).

## Distilling biologically relevant information

The automated processing steps up to this point reduced the data collected per patient 2,000-fold (**Fig. 1a**). However, further curation and prioritization of the information is required to derive interpretable tumor biology. An important challenge is deciding which signals are reliable and which represent technical limitations. To address the latter, protein quantifications based on fewer peptides than expected and quantifications below the dynamic range of the MS signal within the batch were flagged (**Methods, Fig. 3a**). The few missing values for phosphopeptides within a batch were imputed at the limit of the dynamic range, and protein missing values within a batch were flagged as tentative evidence of absence. In addition, for each protein and phosphopeptide, quantification robustness was assessed by the ratio of biological over technical CV (**Fig. 3b, Methods**). This led to their classification as robust (ratio > 2), variable (1 < ratio < 2), unreliable (ratio < 1) or uncharacterized (detected in too few technical replicates).

**Fig. 3:**
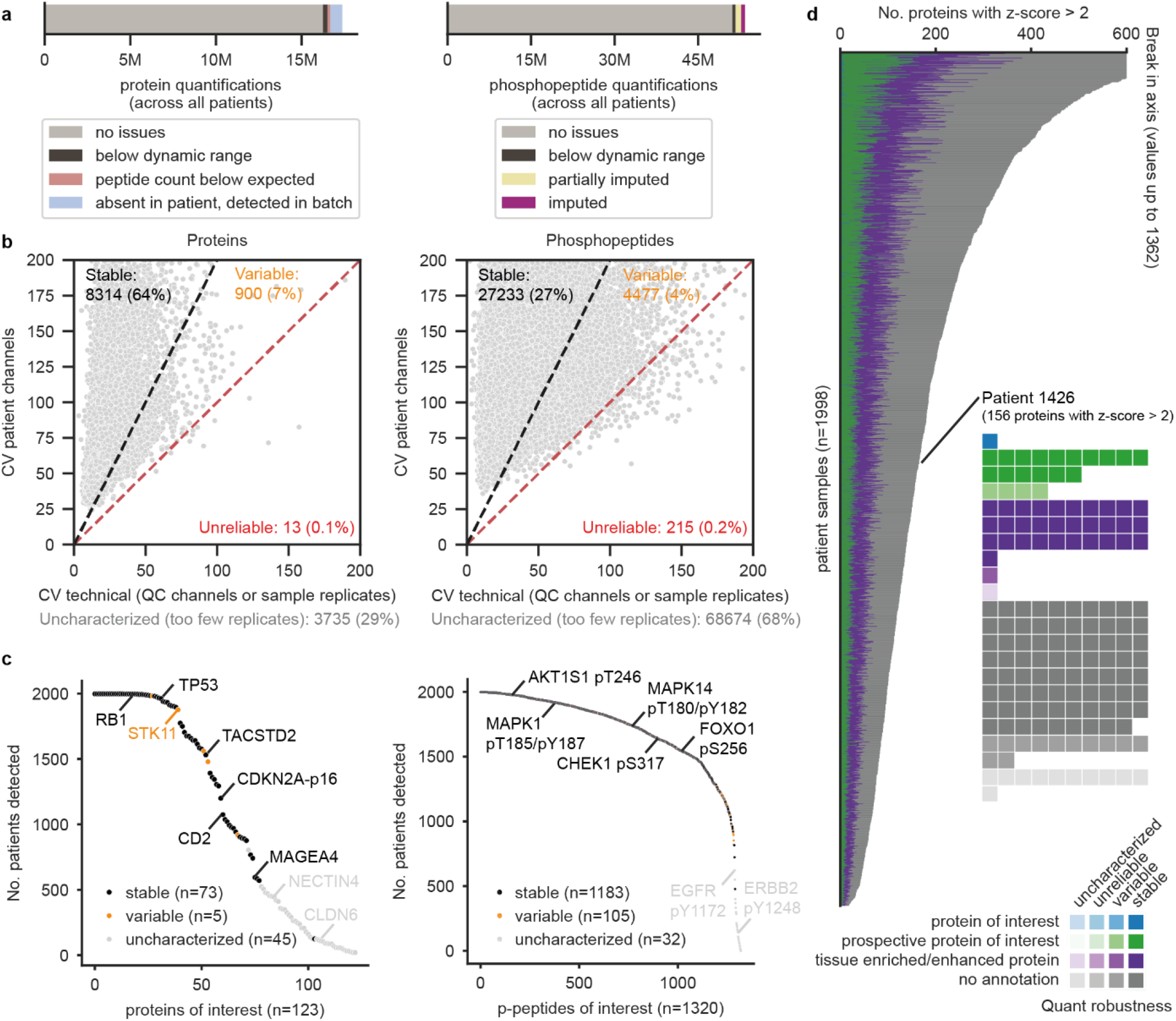
Annotation of quantification robustness and biological relevance of proteins and p-peptides. (a) Flags for potential TMT-induced batch effects in protein and phosphopeptide quantification serve as a warning for the interpretation of such values. (b) Comparison of the biological vs. technical coefficient of variation (CV) of detected proteins and phosphopeptides allows quantification robustness assessment as stable, variable, unreliable, or uncharacterized (detected in too few technical replicates). (c) Quantification robustness assessment by occurrence within the MTB cohort of proteins and phosphopeptides of interest, including oncogenic drivers, tumor suppressor genes, immunotherapy targets, and curated kinase substrates. (d) Number of proteins with a z-score above 2 (≥98th percentile) for each patient that are considered for further investigation. For patient P1426 (inset), these proteins are broken down by biological relevance and quantification robustness (middle right panel).

A straightforward way to prioritize aberrant biological signals is to analyze abundances of known entity biomarkers, tumor antigens, oncogenes, tumor suppressors, drug targets or phospho-sites, annotations of which were obtained from OncoKB, KinHub, ADCdb, PSP and the literature^20–22^ (**Fig. 3c, Extended Data Table 2**). To evaluate if a patient shows abnormal abundance of a protein or phosphopeptide of interest, one might compare the molecular profile to that of nearby healthy tissue^23^. In a precision oncology setting, healthy control samples are not systematically available, and the tissue of origin may be unknown, e.g., for cancers of unknown primary (CUP) or many rare subtypes. Detecting differences in the phosphoproteomes of proliferating cancer cells and fully differentiated (non-proliferating) normal cells does not necessarily imply that they are oncogenic signals. Therefore, we chose all samples contained in this study as a background cohort and computed z-scores for each protein or phosphopeptide for each patient sample relative to the cohort (**Methods**). Z-scores ≥ 2 were typically considered for further investigation (**Fig. 3d),** though clinically relevant results below this threshold were still reported if, e.g., tumor cell content was low. Analyte abundance relative to other patients from the same (or closely related) entity as well as tissue topology (tumor/metastasis site) were also taken into account^10^.

## Tumor proteome activity status (TOPAS) scoring

An exciting potential of patient phosphoproteome data is to infer dysregulation of kinase activity and affected signaling pathways, and several tools have been developed for this purpose (KSEA, INKA, PTM-SEA, KSTAR, ROKAI)^24–28^. These methods strongly depend on reliable kinase-substrate relationship (KSR) annotations but particularly for intracellular kinases (ICKs), KSR databases contain many false positives. This is because the underlying data did not distinguish between direct kinase substrates and phosphorylation mediated by downstream kinase activity^29^. To alleviate these issues, we built a series of TOPAS scores as follows (**Fig. 4a, Methods**). For 18 receptor tyrosine kinases (RTKs), we developed TOPAS-RTK scores by summing up three subscores: 1) kinase protein abundance (overexpression can be a mechanism of overactivity^30, 31^), 2) abundance of (important) kinase activity-indicating phosphorylation (intersection of sites curated from the literature, responding consistently to drug perturbation^29, 32^, and hyperphosphorylated in patients with oncogenic mutations, **Methods**), and 3) total abundance of kinase phosphorylation (**Fig. 4b, Extended Data Table 3**). Adaptor protein phosphorylations by RTKs are not considered in the score because they are not systematically known and the same adaptors are often shared by several RTKs, leading to ambiguity. The 28 developed ICK-TOPAS scores solely relied on the summed abundance of high-quality substrates. These were compiled from coherent behavior in a large-scale dose-dependent phosphoproteomics screen in response to kinase inhibitors recently developed by the authors^29^. Depending on the kinase, ICK-TOPAS scores were based on few or many known substrates (**Fig. 4c**). The robustness of each score varies, as not all proteins or phospho-sites comprising the scores are equally stably detected across patients (**Fig. 4d**). Analogous to phosphopeptide or protein abundance, TOPAS scores are computed as z-scores across the cohort to simplify interpretation.

**Fig. 4:**
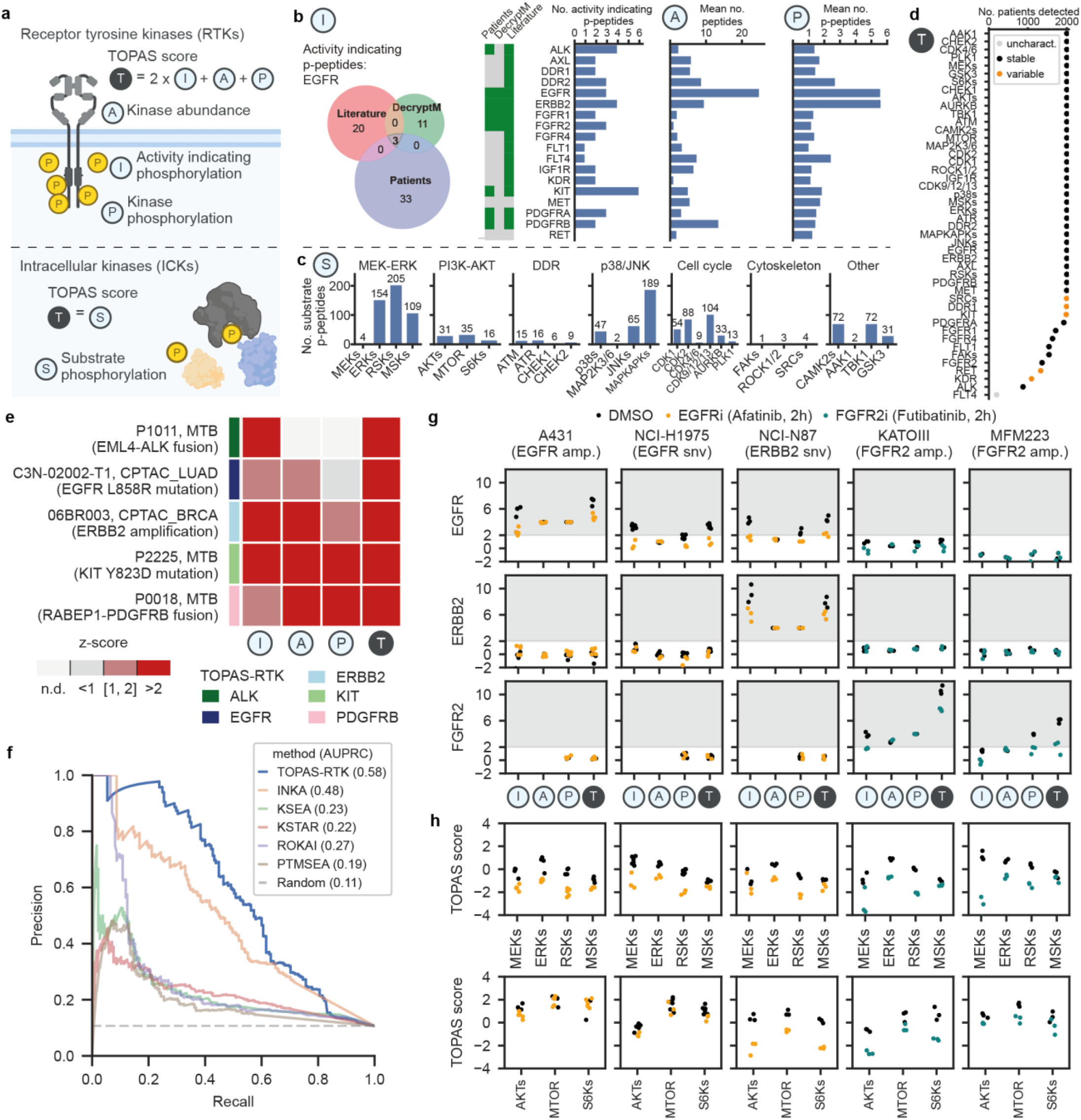
Development and validation of TOPAS scores for detecting aberrant kinase signaling in patients. (a) Schematic representation of the signaling events initiated by a receptor tyrosine kinase (RTK) dimer. The overall TOPAS score (T) combines one (ICKs) or three (RTKs) subscores as indicated. (b) Selection of activity-indicating p-peptides for EGFR using the intersection of literature review, dose-resolved drug-phosphorylation modulation (decryptM), and sites detected in patients of the MTB cohort with oncogenic alterations in EGFR. To the right are the respective numbers used for each RTK as well as the numbers of peptides and p-peptides used for each RTK abundance and phosphorylation subscore. (c) The number of substrate p-peptides used for ICKs. (d) Robustness and occurrence within the MTB cohort of TOPAS-RTK and TOPAS-ICK scores as assessed by technical variation for QC and patient replicates. (e) TOPAS-RTK scores for selected patients in the MTB and CPTAC cohorts with clinically actionable RTK oncogenic driver signals. (f) Precision-recall analysis comparing kinase activity inference methods, evaluated on the full set of patients with clinically actionable RTK oncogenic driver signals. (g) TOPAS-RTK scores (EGFR, ERBB2, FGFR2) for drug perturbation experiments on cell lines with known RTK oncogenic driver signals. Scores without drug treatment are shown as black dots, with drug treatment as colored dots. (h) TOPAS-ICK scores (MEK-ERK, PI3K-AKT-mTOR pathways) for the drug perturbation experiments in (g).

To verify the ability of TOPAS scores to detect dysregulated RTK signaling in patients, we analyzed 94 cases from our own cohort and 79 from the CPTAC cohorts. These patients carry known oncogenic genomic alterations with therapeutic implications in nine RTKs (ALK, EGFR, ERBB2, FGFR1, KIT, MET, PDGFRA, PDGFRB, RET) and were defined to represent ground truth (**Fig. 4e, Methods, Extended Data Table 4**). Although far from perfect, TOPAS-RTK performed better than INKA and both were far superior to several alternatives mentioned above (**Fig. 4f**). The TOPAS sub-scores were of broadly equal performance making TOPAS robust against relying on one metric alone (**Extended Data Fig. 5a**). In addition, results of TOPAS scoring were consistent between patients in our cohort and those in CPTAC (**Extended Data Fig. 5b**). To validate TOPAS scores by experiment, we treated five cell lines with known oncogenic driver alterations in EGFR (A431, NCI-H1975), ERBB2 (NCI-H87) and FGFR2 (KATOIII, MFM223) with afatinib (EGFRi) or futibatinib (FGFR2i) (**Fig. 4g**). Indeed, each cell line showed a significant TOPAS score for its driver kinase, which was reduced upon the respective drug treatment. Extending this analysis to ICK signaling activity in the RAF-MEK-ERK and PI3K-AKT pathways showed that their TOPAS was generally also reduced upon RTK inhibition (**Fig. 4h**). In three cases, the PI3K-AKT pathway was not inactivated upon kinase inhibition, likely due to mechanisms that decouple the RTK from the downstream pathway: A431 (low PTEN levels), NCI-H1975 (PIK3CA mutation), and MCM223 (PIK3CA mutation). As expected, kinase activities in other pathways remained unaffected (**Extended Data Fig. 6**).

## Patient-centric analysis in the TOPAS Portal

To enable molecular oncologists to interrogate and interpret patient phosphoproteomes, we built the TOPAS Portal, an interactive web-based decision support system comprised of several dashboards, each addressing specific aspects of the data (**Fig. 5a**, https://topas-portal.kusterlab.org/). Additionally, patient-specific reports can be downloaded in tabular format for offline investigation and manual annotation of results. Using data from a testicular teratoma patient (P1306) as an example, we illustrate how the portal and reports can be used to prepare for the MTB meeting. First, the overall data quality for the patient and the TMT batch in which the patient was contained is assessed (**Fig. 5b**). This includes checking if i) the summed intensities of peptides and phosphopeptides are in the acceptable range, ii) the QC samples contained in the same batch cluster with the QC samples of all other batches in a principal component analysis (PCA) and iii) other patients in the same batch who are typically from different entities do not cluster together in the PCA. For P1306, all three criteria were met. Failing QC criteria can lead to excluding the patient or the entire batch. Alternatively, one can choose to continue the analysis with consideration of the problematic data quality. In the target identification step, aberrant kinase activity can be explored using TOPAS scores, and aberrant abundance of biomarkers relevant for treatment recommendations, such as tumor antigens, immune checkpoint proteins, immune cell infiltration, or kinase substrate phosphorylation, can be considered (**Fig. 5c**). In our example, EGFR should be considered because of an outstanding TOPAS score. ERBB2 had a TOPAS score just below threshold, indicating potential HER1-HER2-heterodimer signaling. CLDN6 is a noteworthy tumor antigen in this patient because an outstanding response to CLDN6-specific CAR-T cell treatment has been reported in a teratoma patient before^33^. For target verification (**Fig. 5d**), each TOPAS subscore is evaluated individually to check if it supports the conclusion of aberrant kinase activity implied by the overall TOPAS score. This includes looking up phospho-site annotations in PhosphositePlus (PSP) and their reported relevance for kinase activity. For P1306, all TOPAS subscores supported high EGFR activity. Next, the molecular oncologist summarizes the results in a target evidence table (**Fig. 5e**) that lists individual proteins that are clinically actionable, along with the molecular evidence derived from the phosphoproteome data. Potential inclusion of the patient in an ongoing clinical trial is also documented. Results from all four steps are presented in the MTB, supported by plots downloaded from the portal, and discussed alongside the genomics and transcriptomics data collected for the same patient. Final target recommendation is made by accredited clinical oncologists.

**Fig. 5:**
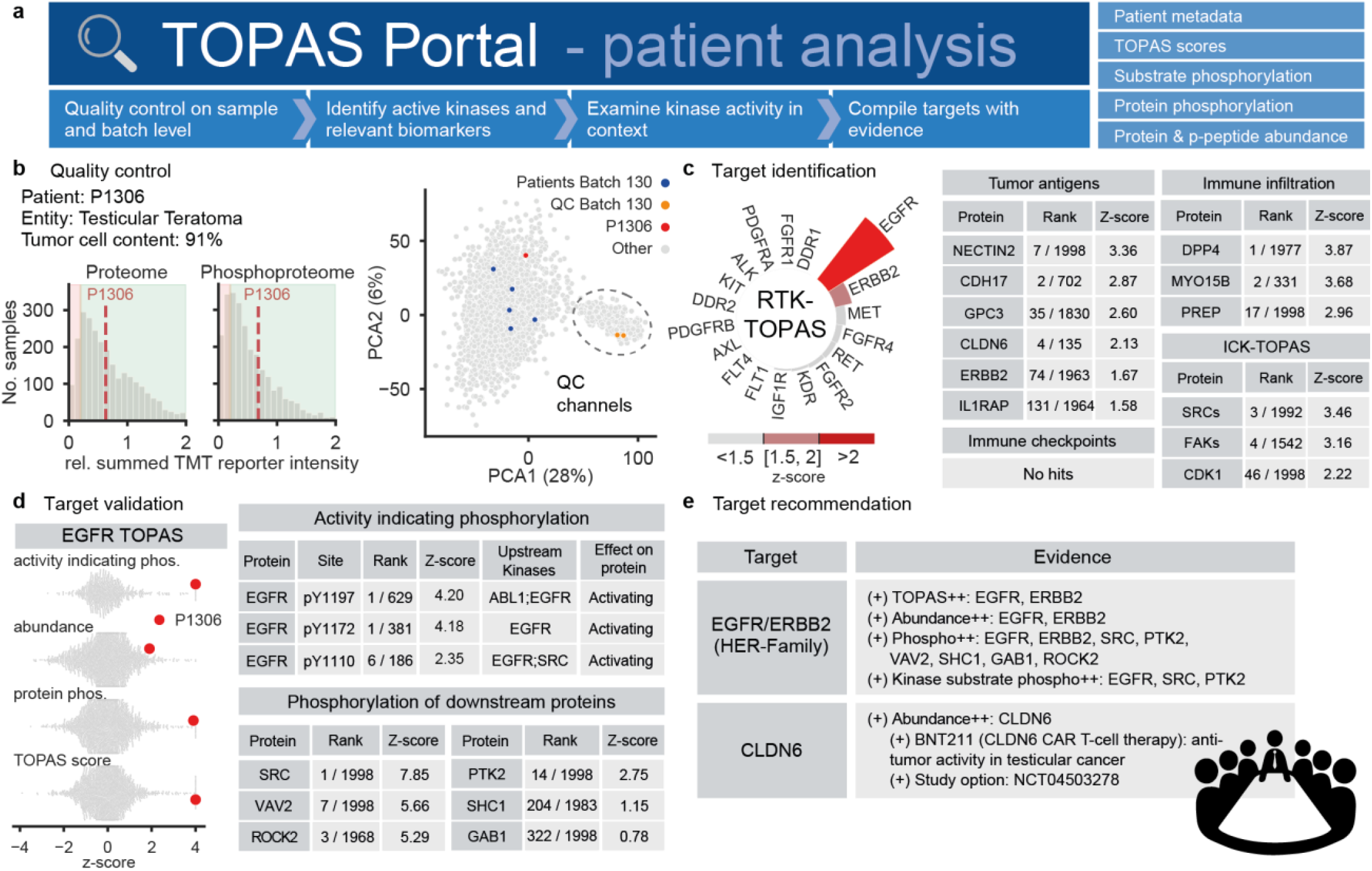
Patient-centric analysis in the TOPAS Portal. (a) The portal consists of dashboards that help address specific questions relevant for the MTB. (b) Quality control: distributions of the summed TMT reporter intensity, relative to the QC channels, of peptides and phosphopeptides for all patients in the cohort, highlighting the patient of interest (P1306, left panel). PCA plot of the proteomes of all patients, highlighting the patient of interest in red, the QC samples contained in the same TMT batch in orange, and the other patients in the same batch in blue (right panel). (c) Target identification: circular plot ranking RTK target proteins by TOPAS score (left panel). Table of proteins and kinase substrates with detected outlier behavior (rank / among number of patients and z-score) relative to the chosen background cohort (right panel). (d) Target validation: swarm plots of TOPAS subscores of all patients (grey) and highlighting the patient of interest in red (left panel). Table of phosphorylation sites with detected outlier behavior (rank / among number of patients and z-score) relative to the chosen background cohort (right panel). (e) Target recommendation: table summarizing the molecular evidence supporting a particular target for discussion in the MTB.

## Cohort-centric analysis in the TOPAS Portal

Molecular data collections on pan-cancer cohorts are highly valuable for discovering common and distinct aspects of tumor biology. In the prospective setting of an MTB, this is particularly true as it may drive therapy decisions. Creating new hypotheses is especially important for rare or otherwise hard-to-treat cancers, a focus of our precision oncology program. The portal offers functionalities to i) detect molecular subgroups within or across entities, ii) compare groups of patients by genomic alteration, iii) detect proteins, kinases, or pathways exhibiting outlier behavior in a group of patients, or iv) discover groups of co-regulated proteins and phospho-sites to name a few (**Fig. 6a**).

**Fig. 6:**
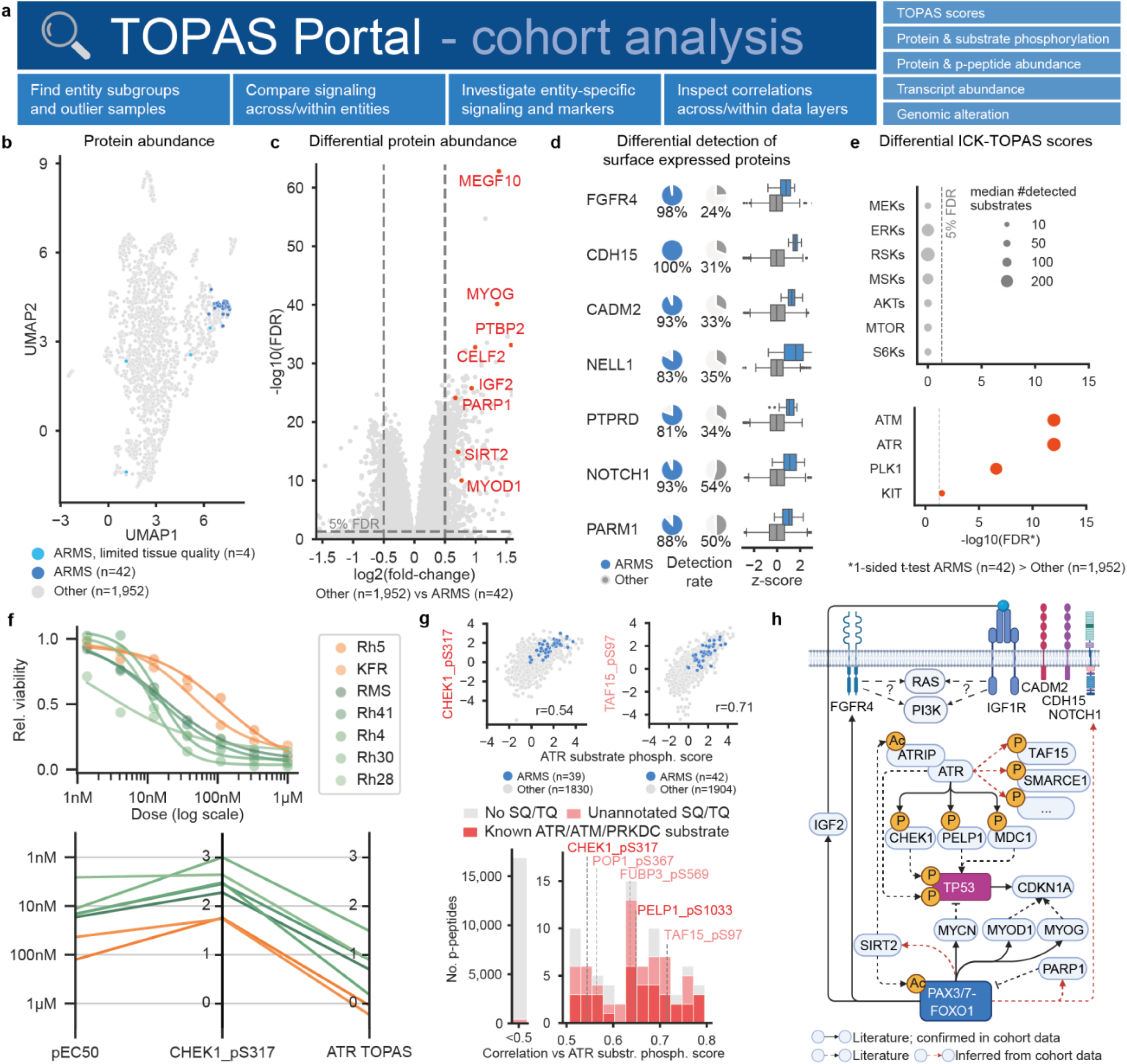
Cohort-centric analysis in the TOPAS Portal. (a) Schematic of the dashboards available in the TOPAS portal. (b) UMAP analysis for protein abundance showing clustering of ARMS samples of high tissue quality in dark blue, four dispersed cases of limited tissue quality in light blue, and all other patients in grey. (c) Volcano plot illustrating differential protein abundance of ARMS vs. all other patient samples. Proteins of interest are highlighted in red. (d) Detection rates and differential abundance analysis of cell surface proteins as potential immunotherapy targets in ARMS (blue) and other patients (grey). (e) Results of differential ICK-TOPAS scores of ARMS patients vs. other cancer entities. The vertical dashed line indicates the 5% false discovery rate (FDR) threshold. (f) Parallel coordinates plots for 7 ARMS cell lines displaying cell viability pEC50 upon ATR inhibition, z-score abundance of the known ATR substrate CHEK1_pS317, and the ATR TOPAS score. (g) Examples of phosphopeptide abundances correlating with ATR substrate phosphorylation scores across all patients for the known ATR substrate CHEK1_pS317 and a novel candidate TAF15_pS97 (upper panel). Number and distribution of such correlations across all frequently observed phosphopeptides (lower panel). (h) Model of potential oncogenic signaling in ARMS based on the literature and augmented with findings from the current study.

Below, we illustrate how the information in the portal can be leveraged and novel biological insights generated using Alveolar Rhabdomyosarcoma (ARMS) patients from the MASTER/INFORM cohort as an example. RMS is very rare but the third most common pediatric solid cancer with an incidence of 4.5 new cases per 1,000,000 children^34^. There are two major subtypes, embryonal and alveolar rhabdomyosarcoma (ERMS, 75%; ARMS, 20% of cases), with ARMS patients having a worse prognosis than ERMS patients (five-year survival of 48% and 73%, respectively), highlighting the need for alternatives to standard treatments^34^. To investigate entity-specific signaling, the portal offers differential expression analysis on every data layer, from protein abundance to TOPAS (sub)scores, and the abundance of individual phosphopeptides. All but four of 46 ARMS patients formed a tight cluster in a UMAP analysis of protein abundance (**Fig. 6b**). Using the metadata annotations provided in the portal, the six outlier cases either had low tumor cell content or failed RNA library preparation, usually indicating reduced overall sample quality. These samples were excluded from subsequent analyses. About 80% of ARMS patients have a chromosomal translocation leading to a PAX3/7-FOXO1 fusion^35^. This fusion promotes the transcription of several genes, including the RTKs FGFR4, IGF1R, ALK, and MET as well as the transcription factors MYOD1 and MYOG^35–37^. Differential protein abundance analysis revealed 509 significant associations (t-test) for ARMS vs. all other patients (**Fig. 6c**). These included many tissue-enriched proteins as classified by the human protein atlas (skeletal muscle: 4 tissue enriched, 50 group enriched, 34 tissue enhanced)^38^ but also several PAX3/7-FOXO1 target genes (**Extended Data Fig. 7a**).

A similar analysis for cell surface proteins^39^ revealed 10 proteins (Fisher’s exact test) including the cell adhesion proteins CDH15 and CADM2^40^ as well as two receptors, NOTCH1^41^ and FGFR4 (**Fig. 6d**). Interestingly, elevated expression of FGFR4 did not result in overactivity of downstream kinases in the PI3K/AKT or RAS/MAPK pathways (**Fig. 6e**). While FGFR4 inhibition has been shown to provoke a response in RMS cell lines^42^, treatment of xenograft mouse models was ineffective^43^. We, therefore, hypothesize that FGFR4 expression does not appear to drive the growth of ARMS tumors. Instead, ICK-TOPAS scoring identified four potentially overactive kinases in ARMS patients that may represent therapeutic vulnerabilities (**Fig. 6e**). The two most significant hits were ATM and ATR, both members of the PIKK family that share substrates bearing the SQ/TQ motif. The observed high ATR substrate scores provided a molecular rationale for the reported efficacy of ATR inhibitors in ARMS cell lines^44^. To investigate if ATR/ATM TOPAS scores are predictive of reduced cell viability upon ATR inhibition, we measured phosphoproteome profiles for 7 ARMS cell lines. The well-known ATR substrate CHEK1_pS317 had a z-score above two for the five cell lines with strong response to ATR inhibition by Elimusertib (EC50 < 20nM), whereas the two cell lines with weak response (EC50 > 40 nM) had elevated but non-significant z-scores. Four cell lines with strong responses had ATR TOPAS scores > 0.5, whereas the two cell lines with the weakest response had scores below zero (**Fig. 6f**).

The high diversity in ATR activity across the pan-cancer cohort also provided an opportunity to find new ATR/ATM substrates. This was achieved by correlating the abundance of SQ/TQ phosphopeptides that have not yet been annotated as ATR/ATM substrates with the ATR TOPAS scores of patients (**Fig. 6g; Extended Data Fig. 7b**). This approach detected well-known substrates such as CHEK1_pS317 and PELP1_pS1033 but also added 29 potential new ones. Among them are TAF15_pS97, FUBP3_pS569, and POP1_pS367, all of which responded to ATR inhibition by Elimusertib in our recent study in PDAC cell lines^45^.

Another interesting observation of ARMS tumor biology derived from the cohort analysis was an elevated abundance of PARP1 (**Extended Data Fig. 7a**). This rationalizes why combining ATR and PARP inhibitors works synergistically in some ARMS cell lines^44^. Moreover, elevated levels of the deacetylase SIRT2 provided a reasonable hypothesis for the high ATR activity in ARMS because SIRT2 is known to drive ATR checkpoint activation by deacetylation of ATRIP^46^. SIRT2 also deacetylates FOXO1, resulting in nuclear import and increased transcriptional activity of FOXO1^47^. This results in a feedback loop in which FOXO1 targets the promoter region of SIRT1^48^. Although no such feedback loop has been described for SIRT2 yet, its promoter region contains a binding site for FOX transcription factors^49^. This further suggests that SIRT2 inhibition may be even more effective than ATR inhibition^50^. The model shown in **Fig. 6h** derived from the cohort analysis generated several new hypotheses regarding aberrant signaling in ARMS and identified ATR, PARP1 and SIRT2 as actionable targets, potentially paving the way for new targeted therapies.

## Discussion

The TOPAS platform was developed to assist scientists and clinicians in gaining therapeutically actionable insights from analyzing the phosphoproteomes of individual cancer patients or cohorts in a prospective MTB setting. Such data analysis pipelines for genomics and transcriptomics^51, 52^ as well as clinical decision support systems (CDSS) for genomics^53–55^ have been available to MTBs for some time, but no comparable tools existed for phosphoproteome data. While the CPTAC data analysis pipeline (CDAP)^56^, paired with PANOPLY^57^, a cloud platform offering analysis tools for cohort-wide investigations, follows a unified approach, it is currently not possible to analyze or contextualize the profiles of individual patients. The same is true for the Clinical Knowledge Graph (CKG), which integrates data processing, functional annotation, and visualization within a knowledge graph database^58^. In addition, all available tools today have only been used for retrospective cohort analysis. To derive tailored options for each patient in need of therapy, independent of whether it is a common, well-studied, or a rare cancer subtype, or even a cancer of unknown primary, retrospective cohort studies are insufficient.

The TOPAS platform specifically addresses this gap and is the first tool with demonstrated ability to analyze the phosphoproteomes of a large number of prospective MTB cases. While the platform is computationally efficient, the main current bottleneck is the time needed by molecular oncologists to interpret and understand the data for each case well enough to arrive at a treatment recommendation. This can be challenging because it is often unclear which level of over- or underactivity of a signaling component (protein, phospho-site, kinase, pathway, etc.) may constitute an actionable target in the complex biological and disease journey context of this individual patient. The simple approach used here of prioritizing parameters with a z-score of ≥2 carries a risk for both false positives and negatives. In addition, the full potential of the TOPAS scoring concept is currently underutilized because of the lack of comprehensive, high-confidence, and cancer-context-aware kinase-substrate relationships. Recent computational advances in kinase motif analysis^59^ and large-scale phosphoproteome profiling of drug perturbation experiments^29, 32^ can be expected, at least in part, to alleviate this issue.

The TOPAS platform is under continuous development and we are happy to help others set it up in their MTB environment. In the future, visualizations that overlay the signaling components for each patient directly onto curated pathway diagrams^60^ will become available. Additionally, we are collecting training data from manual evaluations of dysregulated signaling pathways in patients to develop machine learning models that can provide faster decision support. By close, multidisciplinary collaboration, we anticipate that phosphoproteome profiling alongside the open-source TOPAS analysis platform has the potential to become a widely used component in precision medicine programs around the globe.

## Acknowledgements

We thank all members of the TU Munich team for high-quality technical and intellectual support. This work was supported by the European Research Council (ERC) grant 833710 (TOPAS), the German Federal Ministry of Education and Research (BMBF), grants 01KD2207F (HEROES-AYA), 01KD2206C (SATURN-3), 161L0214A (CLINSPECT-M), 03LW0243K (CLINSPECT-M-2), 031L0305A (DROP2AI) and 031L0168 (DIAS), the German Research Foundation (DFG) grant 537476536 (PhoSAIC). The MASTER program is supported by the NCT Overarching Clinical Translational Trial Program, the NCT Heidelberg Molecular Precision Oncology Program, and DKTK.

## Author contributions

M.T., A. Schneider and B.K. conceived and designed the study. C.B.J., A. Sakhteman, F.H., A. Schneider and M.T. performed (phospho)proteome data processing and analyses. C.B.J., A. Sakhteman, F.H., J.W. and M.T. wrote software/code. F.P.B., J.H., D.H. and S.F. provided resources. A. Schneider, M.V.T., P.H. and C.S. performed data curation on MTB recommendations and/or clinical data. A.G.H. provided cell line material and phenotypic data of alveolar rhabdomyosarcoma cell lines. M.T., S.F. and B.K. supervised the study. C.B.J., A. Schneider, B.K. and M.T. wrote the manuscript. All authors reviewed and edited the manuscript.

## Competing interests

BK is a non-operational co-founder of OmicScouts and MSAID and has or has had an ownership, consulting or advisory role and/or received honoraria, research funding, and/or travel/accommodation expenses funding from AstraZeneca, BASF, Bayer, Biogen, Boehringer Ingelheim, Bruker, Covant Tx, Dunad Tx, EQT Partners, IBM, Merck KGaA, MSAID, MSD, Nested Tx, OmicScouts, Roche, SAP and ThermoFisher Scientific.

S.F. has or has had a consulting or advisory role and/or received honoraria, research funding, and/or travel/accommodation expenses funding from Illumina and Oxford Nanopore Technologies, outside the submitted work.

## Methods

### Data availability

For the MTB cohort, patient raw phosphoproteome mass spectrometry data, peptide and protein identification and quantification data provided by MaxQuant, as well as TOPAS pipeline results are available from the German Human Genome-Phenome Archive (GHGA, study identifier pending). Because this data contains patient-derived information, controlled access is required and can be granted upon request. Raw phosphoproteome mass spectrometry data files for three published studies provided by CPTAC^1–3^ were downloaded from the CPTAC data portal in May 2021 (https://cptac-data-portal.georgetown.edu/datasets) utilizing the IBM Aspera client. CPTAC peptide and protein identification and quantification data provided by MaxQuant, as well as TOPAS pipeline results are available from PRIDE with the project accession number (PXD061316).

Reviewers can access the dataset by logging into the PRIDE website using the following account details:

Username

Password: eyGZ4OHWnDsv

Raw files for all 44 sarcoma samples from the TCPCA cohort^4^ were downloaded from PRIDE with the project accession number PXD054790 using the Python package pridepy v0.0.16.

### Peptide and protein identification and quantification by MaxQuant

For peptide and phophopeptide identification and quantification of the MTB and CPTAC cohorts, raw mass spectrometry files were searched with MaxQuant (v.1.6.12.0 for Batch 1-229 and all CPTAC datasets, v2.1.3.0 for Batch 230-336) against the human SwissProt+TrEMBL database (97,086 protein sequences including isoforms, downloaded Nov 2020) and the common contaminants database from MaxQuant^5^. To distinguish CDKN2A p14 and p16 splice variants, CDKN2A entries had their gene name extended by the respective splice variant names CDKN2A-p14 and CDKN2A-p16. The proteome and the phosphoproteome datasets were searched separately and per TMT batch. A custom automated pipeline was set up in Python to start a MaxQuant search as soon as all raw files for a batch were measured.

Default parameters of MaxQuant were used with the following changes. Searches were performed using the ‘MS3 reporter ion’ experiment type and ‘TMT11plex’ labels. Isotope impurity correction factors provided by the manufacturer were used for each TMT lot. Searches included up to three missed cleavage sites, a minimum peptide length of 6, maximum peptide mass of 6,000 and a fragment ion tolerance of 0.4 Da (ITMS). PSM-level FDR was set to 1% and protein level to 100% (for protein level FDR control see below). For the phosphoproteome, phosphorylation of serine, threonine, and tyrosine was set as variable modification. CPTAC data were searched with the same parameters except the experiment type was set to ‘MS2 reporter ion’.

### Peptide and protein identification and quantification by FragPipe

For peptide and phophopeptide identification and quantification of the TCPCA cohort, raw mass spectrometry files were searched with FragPipe v23.1^6^ and quantified with DIA-NN v2.3.2^7^ against the same SwissProt+TrEMBL database as described above for MaxQuant. Default parameters for the DIA workflow in FragPipe were used with the following changes: diann.cmd-opts=--report-decoys, msfragger.allowed_missed_cleavage_1=2, msfragger.allowed_missed_cleavage_2=2, phi-report.dont-use-prot-proph-file=true, phi-report.filter=--prot 1.0 --pep 1.0 --ion 1.0, phi-report.print-decoys=true.

### TOPAS pipeline

#### Software and performance

The TOPAS pipeline was implemented in Python 3.9.13 and is available at https://github.com/kusterlab/topas-pipeline. A Docker image is automatically created with every git commit. The pipeline performs the following steps (outlined in further detail below): quality control, MS2 identification transfer, normalization and imputation, protein grouping (proteome only), protein or phosphopeptide quantification, fold-change and z-score calculation, functional annotation, TOPAS scoring and report creation. System and unit tests were implemented to ensure stability and maintainability. The pipeline is run in a Docker container by passing it a configuration file in TOML format with input/output paths and settings for the pipeline run. Compute-intensive steps of the pipeline were optimized to use multiple cores and a pipeline run of ∼2,000 patients is completed in ∼24 hours using ∼250GB of RAM. The pipeline is routinely run on a Linux server with 64 cores and 512GB of RAM which can run several pipeline runs in parallel if needed.

#### Quality control

Batches with TMT labeling efficiency <90% (calculated by ‘summed intensity TMT-labelled peptides’/‘summed intensity all peptides’) or mean IMAC enrichment efficiency <80% (calculated by ‘summed intensity phosphopeptides’/‘summed intensity all peptides’) over the 12 fractions were excluded. Furthermore, samples for which the ‘summed TMT reporter intensity’ was <20% (threshold chosen empirically) of the mean ‘summed intensity of TMT reporter intensity’ of both reference channels were assigned a “QC failed” label and excluded from further analyses. Additionally, samples were also excluded from further analyses if the genomics or transcriptomics data detected contaminations or if the sample identity could not be ascertained (termed excluded by ‘other’). Samples with a tumor cell content <20%, severe DNA degradation, documented long cold ischemia time (>30 min) or RNA library preparation failure were assigned a ‘limited tissue quality’ label.

#### Missing data reduction between patient (TMT) batches

To improve data completeness, MQ output files from all TMT batches were submitted together to SIMSI-Transfer version 0.6.3^8^ to transfer MS2 identifications. The following settings were used: “--tmt_ms_level ms3 --stringencies 10 --tmt_requantify --maximum_pep 1 --num_threads 8”. For the CPTAC dataset, the setting “--tmt_ms_level ms2” was used instead.

#### Normalization and imputation

For the MTB cohort, several normalization steps were performed to correct for differences in protein input amounts and batch effects. A graphical depiction is also given in **Extended Data Fig. 3a**. Within each batch, MS3 intensities were median centered using peptides detected in ≥4 TMT channels. Across batches, MS1 intensities were median centered using peptides detected in >70% of batches. For precursors for which MaxQuant did not report an MS1 intensity but an MS3 signal was measured in one of the QC channels 10 or 11 (5% of peptides and 12% of phosphopeptides), an MS1 intensity was imputed as

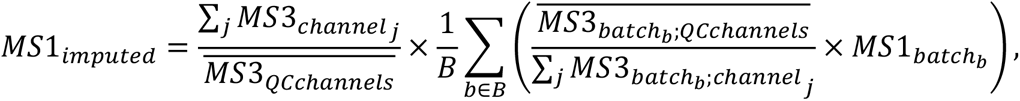

where *B* is the set of batches where an MS1 intensity was reported by MaxQuant. This normalization is based on the assumption that the contribution of the QC channels to the MS1 signal (the term in parenthesis in the summation) is equal in all batches and that is used as a correction factor for the batches where no MS1 intensity was reported.

Within each batch, MS3 intensities were transformed into quasi-LFQ^9^ intensities using the MS1 intensity as

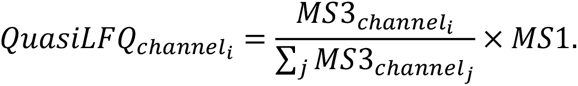

For the CPTAC cohorts, a different approach was necessary because the MS1 peptide chromatograms were insufficiently reproducible between batches. Therefore, within each batch, MS3 intensities were median centered using peptides detected in ≥4 TMT channels. Next, ratios of the MS2 intensities to the common reference channel were taken and transformed into quasi-LFQ intensities as

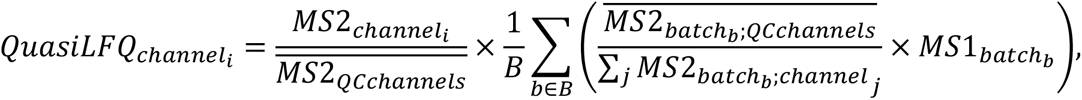

where *B* is again the set of batches where an MS1 intensity was reported by MaxQuant. The term on the right is the average contribution of the QC channels to the MS1 signal. This normalization was applied to each of the three entities (BRCA, LUAD, UCEC) separately.

For the TCPCA cohort, the non-normalized MS1 intensities (“Ms1.Area” column) reported by DIA-NN were used as peptide intensities.

#### Phosphopeptide normalization and imputation (phosphoproteome only)

In addition to the general normalization and imputation strategies described above, additional normalization and imputation steps were performed for phosphopeptides. For phosphopeptides detected in at least 1 QC channel for >67% of batches, intensities were bridge normalized using the QC channels as bridge channels. Finally, for each phosphopeptide, intensities were median centered across different QC lots using the median patient intensity per QC lot. If a phosphopeptide precursor ion was not detected for a channel but was detected in the batch, it was imputed with either i) 1/100^th^ of the maximum intensity or ii) the minimum intensity for the peptide within its batch, whichever of the two was lower. This imputation was based on the fact that ratios smaller than 1/100^th^ can typically not be reliably measured by TMT.

#### Protein grouping and quantification (proteome only)

Protein grouping, 1% protein-level FDR control and MaxLFQ quantification of all TMT batches together was performed on gene level using the picked protein group FDR package v0.7.6^10^. The fasta file used in the MaxQuant database search was used as input and the following arguments were used: “--gene_level --enzyme trypsinp --min-length 6 --cleavages 3 --lfq_min_peptide_ratios 1 --methods picked_protein_group_mq_input --lfq_stabilize_large_ratios”.

#### Phosphopeptide grouping and quantification (phosphoproteome only)

For each phosphopeptide, precursor intensities were summed up across charge states and fractions. To reduce missing values due to ambiguous localization, phosphopeptides with the same naked sequence and number of phosphorylated residues were grouped if each phosphorylated residue was within a ±2 residue window. The most prevalent phosphopeptide within the group was defined as the representative modified sequence. If multiple precursors were obtained for the same group in a single sample, the intensities were summed. If all precursor intensities in a group were imputed, the quantification was flagged as “imputed”, if imputed and non-imputed precursor intensities were combined, the quantification was flagged as “partially imputed”. For simplicity, these groups are referred to as phosphopeptides throughout the manuscript and the remainder of the methods section.

#### Caution flags for quantification values

Besides the “imputed” and “partially imputed” flags described above, three additional flags were employed. First, false identification and inaccurate quantification of peptides can distort protein quantifications, especially when based on few peptides. To detect such problematic cases, we performed a linear regression for each protein, modeling its intensity as a function of the detected peptide count *C* across samples. Protein quantification with a residual of 2 × 1.1*^C^*^−1^ standard deviations above the predicted protein intensity were flagged as “pepcount_OOR”. The multiplicative factor 1.1*^C^*^−1^ was empirically determined to produce more lenient thresholds as the peptide count increases. Second, quantification values below 1/100^th^ of the maximum quantification value in the batch were flagged as “quan_OOR”, again informed by the observation that ratios smaller than that can typically not be reliably measured by TMT. Third, if a protein was not detected in a channel but detected in one or more other channels in that batch, the missing quantification value was flagged as “detected in batch” to indicate a potential true absence.

#### Fold change and z-score calculation

Protein and phosphopeptides intensities were log10-transformed. For each protein and phosphopeptide, the rank, z-score, and fold change were calculated relative to the cohort. For the z-score and fold change, the median was used instead of the mean and the abundance for the patient itself was excluded from the computation of the median and standard deviation.

#### Functional annotation

Cancer gene annotations were downloaded from OncoKB (https://www.oncokb.org/cancer-genes, accessed 22 Aug 2024). Kinase gene annotations were downloaded from KinHub (http://www.kinhub.org/kinases.html, accessed 17 Jan 2024). ADC targets were downloaded from ADCdb (http://adcdb.idrblab.net/, accessed 19 Aug 2024). Immunotherapy targets were collected from the literature^11–13^. The full list of annotations can be found in Extended Data Table 1. Phosphopeptides were annotated using the Python package *psite_annotation*v0.5.3 (https://github.com/kusterlab/psite_annotation) using the following files downloaded from PhosphoSitePlus (https://www.phosphosite.org/staticDownloads, accessed January 13th 2024): Phosphosite_seq.fasta, Kinase_Substrate_Dataset, Phosphorylation_site_dataset, Regulatory_sites. RNA tissue specificity classifications were downloaded from the Human Protein Atlas (https://www.proteinatlas.org/download/tsv/rna_tissue_hpa.tsv.zip, accessed May 11^th^ 2026). Gene annotations classified as “Tissue enhanced” or “Tissue specific” were retained together with the corresponding tissue(s).

#### CPTAC analysis

The three CPTAC cohorts were preprocessed together by the TOPAS pipeline, where each cohort was assigned its own QC lot number.

### Tumor proteome activity status (TOPAS) scores

#### Kinase abundance score

The kinase abundance score of a patient is the z-score of that patients’ kinase abundance computed relative to the background cohort consisting of all other patients, and is computed independently for every kinase.

#### Activity indicating phosphorylation score

For the activity indicating phosphorylation score a set of phosphopeptides (typically, well-known autophosphorylation sites) are selected as the intersection of sets of phosphopeptides i) uniquely attributed as substrate of the RTK according to PhosphoSitePlus, ii) regulated by a drug selectively targeting the RTK in a decryptM experiment^14, 15^ and iii) differentially abundant between patients with a known oncogenic alteration in the RTK versus the rest of the MTB cohort. If one or two of these sets were unavailable for an RTK, an intersection was done with the remaining sets. For each patient, the log10 fold change is computed relative to the background cohort for each phosphopeptide included in the set. To arrive at the protein phosphorylation scores, the sum of log10 fold changes are z-scored across all patients, resulting in standardized scores that can be compared to each other.

#### Protein phosphorylation score

For the protein phosphorylation score, all phosphopeptides on the protein are considered, excluding phosphopeptides shared with other proteins. As with activity indicating phosphorylation scores, the protein phosphorylation scores is the sum of log10 fold changes, z-scored across all patients.

#### Substrate phosphorylation score

For substrate phosphorylation scores, we first removed the protein expression contribution for each phosphopeptide by fitting a log-log-linear model with a bounded slope between 0 and 1, and subtracted the regressed linear protein expression effect from the phosphosite abundance. For the substrate phosphorylation score, all phosphopeptides annotated as a substrate of the kinase according to *Bayer et al.*^15^, detected in at least 1 QC channel for 67% of batches (i.e. those for which bridge normalization was performed) are considered. For MEKs, MAP2K3/6, CHEK1 and AAK1, only the seed sites were used as kinase substrates. As with activity indicating phosphorylation scores, the substrate phosphorylation scores is the sum of log10 fold changes, z-scored across all patients.

#### TOPAS scores

The TOPAS-RTK scoring consists of three subscores: kinase abundance, activity indicating phosphorylation and protein phosphorylation. The TOPAS-ICK scoring consists of a single subscore: substrate phosphorylation. For each TOPAS score, the subscores are summed up with the activity indicating phosphorylation subscore weighted doubly. Finally, the summed scores are again z-scored across all patients to arrive at a standardized TOPAS score for each of the 18 RTKs and 28 ICKs for each patient.

### TOPAS portal

#### Data portal

The TOPAS portal (https://github.com/kusterlab/topas-portal) was implemented in Python v3.9.12, Flask v2.3.3 (backend), and Vue v2.6.14 (frontend). The backend supports both an in-memory (default) and a postgreSQL database for the data matrices. Docker images were created for both the frontend and backend which are deployed together using docker-compose. The TOPAS portal supports multi-omics patient data (proteomics, transcriptomics and genomics) and provides access to all the different data modalities (protein, phosphopeptide and transcript abundance as well as TOPAS (sub)scores). It offers differential abundance (t-test, ANOVA, multiple testing correction), cluster (PPCA, UMAP, silhouette scores) and correlation analysis (Pearson correlation). All analyses can be performed on or between subgroups of the cohort based on the provided metadata (e.g. cancer entity, genomic alteration, batch, etc.) or by uploading lists of sample identifiers. This includes the option to compute ranks, z-scores and fold changes relative to, e.g. other patients from the same (or similar) entity or from the same tissue topology (in case of metastatic samples). Visualizations include swarm, volcano, kernel density estimation, correlation and bar plots as well as histograms, heatmaps and Venn diagrams. Endpoint validation and unit tests were implemented to safeguard stability.

#### Report creation

For each patient, a report in Excel format is created consisting of 4 tabs: Summary (containing TOPAS scores and proteins of interest), Proteome, Phosphoproteome, and Protein phosphorylation score. Each of the tabs contains the abundance/score, z-score, rank within cohort, rank within batch, fold-change and, if applicable, functional annotation columns and if the abundance/score data was based on imputed data for this patient.

### Cell culture and drug perturbation assays

#### TOPAS score cell line validation

A431, NCI-H1975 and NCI-N87 were cultured in RPMI-1640 (Gibco, A10491) supplemented with 10% FBS (Pan Biotech). KATOIII was cultured in IMDM supplemented with 20% FBS (Pan Biotech) and MFM-223 was cultured in DMEM supplemented with 15% FBS and 1% 100x ITS. Cells were cultured in a humidified cell incubator at 37 °C and 5 % CO2 and regularly checked for mycoplasma contamination. For perturbation experiments, cells were seeded in 10 cm² dishes and grown to 80% confluence within 48 h. Cell culture medium was replaced 24 h after seeding. A431, NCI-H1975 and NCI-N87 cells were treated with 100 nM Afatinib, 1 μM Lapatinib, 10 μM Osimertinib, or DMSO vehicle for 4h (NCI-H1975 and NCI-N87) in duplicates or with 500 nM Afatinib or DMSO vehicle for 2h (A431) in triplicates. KATOIII and MFG-223 cells were treated with 100 nM Futibatinib, or DMSO vehicle for 2h in triplicates. Cells were washed twice with PBS and lysed in SDS lysis buffer (2% SDS in 40 mM Tris-HCl, pH 7.6). For DNA hydrolysis, samples were heated at 95 °C for 10 min and incubated with 2% trifluoroacetic acid (TFA) for 1 min, before the reaction was quenched using 4% N-methylmorpholine (NMM). Lysate was cleared by centrifugation at 10,000 × g for 10 min and protein concentration was determined using the Pierce BCA Protein Assay Kit (Thermo Scientific) and cells were further processed in the same way as patient’s tumor tissue specimen in separate cell line TMT-batches as described in our accompanying manuscript^16^.

#### Alveolar Rhabdomyosarcoma

Alveolar rhabdomyosarcoma cell lines (Rh4, Rh5, Rh30, Rh41, RMS, KFR) were cultured and harvested as previously described^17^. Cells were further processed in the same way as patient’s tumor tissue specimen in separate cell line TMT-batches as described in our accompanying manuscript^16^. Cell viability data were taken from *Dorado García et al.*^17^ and re-processed using CurveCurator v0.6.0^18^.

### Data analysis

#### Comparisons to data matrices before normalization

The normalized data matrix was compared to a pre-normalized data matrix computed by skipping both the within (MS3) and across (MS1) batch median centering but including calculation of quasi-LFQ intensities. All other steps in the TOPAS pipeline were executed identically.

#### Comparison to summed intensity protein quantification

Summed intensity protein quantification was performed on gene level using the picked protein group FDR package v0.7.6^10^ using the same settings as described for MaxLFQ in the section “Protein grouping and quantification”. All other steps in the TOPAS pipeline were executed identically.

#### Comparison to median centering without occurrence filter

The minimum occurrence filter on proteins or phosphopeptides used for median centering within and across TMT batches was set from 70% to 0%. All other steps in the TOPAS pipeline were executed identically.

#### Quantitative precision analysis

Non-log-transformed intensities were used to compute the coefficient of variation (CV) as a metric for quantitative precision for all proteins and phosphopeptides found in >70% of samples. Cumulative density plots for CV were created for sample replicates and QC channels across batches. For sample replicate CV calculation, replicate samples for P0291 measured in three batches (Batch 29, 30, 31) were used. For both TMT Channel 1 (patient samples) and 9 (QC samples), CVs were computed across batches for the corresponding lot of QC samples (batches 1-72).

#### Quantification robustness analysis

For each protein and phosphopeptide, CV was calculated on non-log-transformed intensities as CV = (1 + 1/4n) × SD/mean × 100, with the (1 + 1/4n) term correcting for small-sample bias. Technical CV was computed from QC-channel replicates or from technical replicates of the same patient sample. Technical CV was the median across TMT batches/lots (if detected in ≥70% of TMT batches) or patient replicate sets (if detected in ≥5 patients), whichever of the two had higher detection coverage. Biological CV was simply computed across all patient samples (if detected in ≥20 patients). For TOPAS (sub)scores, standard deviations were calculated instead of CVs using the same criteria as described above for CVs. Biological-to-technical CV or standard deviation ratios were classified as “unreliable” (ratio <1), “variable” (ratio 1–2), “stable” (ratio >2), or “uncharacterized” (detected in too few QC channels and patient replicates).

#### Standard error of the mean reduction analysis

For each protein and phosphopeptide, log-transformed intensities were used to compute the standard error of the mean for each entity with at least 10 patients. The reduction in standard error of the mean (SEM) was calculated as 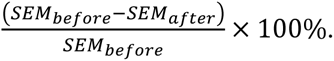 Finally, the median reduction in SEM was computed across all protein-entity or phosphopeptide-entity pairs.

#### Biomarker differential expression

Known protein biomarkers for the Solitary Fibrous Tumor (SFT) and known protein and putative phosphopeptide biomarkers for Ewing Sarcoma (ES) entities were selected based on literature research. *P* values were calculated between the protein or phosphopeptide abundances of patients within the entity against the rest of the cohort using the two-sided independent t-test function *ttest_ind()* of scipy v1.11.4.

#### UMAP and PPCA

Proteins found in >50% of samples were retained. Missing values were imputed using the PPCA method included in scikit-learn v1.3.0. UMAPs were plotted from the imputed data matrices using 10 neighbors and 1,000 epochs.

#### Kinase activity scoring comparison

INKA^19^, KSEA^20^, RoKAI^21^, PTM-SEA^22^ and KSTAR^23^ enrichments were performed using the enrichment-server v0.1.3 of PTMNavigator^24^. For INKA, the intensities were used as input for each phosphopeptide. For KSEA, RoKAI and PTM-SEA, the log2 fold changes of all patients relative to the median were used as input for each phosphopeptide. For KSTAR, only log2 fold change > 1.0 were used as input for each phosphopeptide. For the patient analysis, TOPAS-RTK scores were extracted for patients with at least one oncogenic RTK alteration with a therapeutic evidence level according to OncoKB (accessed 18 May 2026). The oncogenic altered RTKs for each patient were considered as real positives, whereas all other RTKs for the patient were considered as real negatives. For the MCC and confusion matrix analyses, the following thresholds were employed: z-score > 2 for TOPAS (sub)scores, summed intensity > 8.0 for INKA, and FDR < 5% for KSEA, RoKAI, PTM-SEA and KSTAR.

#### Differential protein abundance analysis

For each protein, the *p* value was calculated between the protein abundances of ARMS samples (excluding the six samples with degradation or low tumor cell content issues) and all other samples in the cohort using the independent t-test function *ttest_ind()* of scipy v1.11.4. Benjamini-Hochberg multiple testing correction was performed with the *fdrcorrection()* method of statsmodels v0.14.2. Proteins were classified as significant if FDR <0.05 and log2(fold change) >0.5. Classification of tissue specific proteins was done using the 921 elevated genes for skeletal muscle reported in the Human Protein Atlas^25^ (https://www.proteinatlas.org/humanproteom/tissue/skeletal+muscle, accessed 19 Sep 2024).

#### Differential detection of surface expressed proteins

Surface expressed proteins were extracted from Supplementary Table S2A of the Cell Surface Protein Atlas publication^26^. Only proteins with the CSPA category of “1 – high confidence” were retained. For each protein, the *p* value was calculated between the presence of the protein in ARMS samples (excluding the six samples with degradation or low tumor cell content issues) and all other samples in the cohort using the fisher exact test *hypergeom.sf()* of scipy.stats v1.11.4. Benjamini-Hochberg multiple testing correction was performed with the *fdrcorrection()* method of statsmodels v0.14.2. Proteins were classified as significant if FDR <0.01, log2(fold change) >0.4 and the protein was detected in more than 30 of the 42 ARMS samples.

#### Differential kinase substrate phosphorylation analysis

For each kinase, a *p* value was calculated between the TOPAS-ICK scores of ARMS samples (excluding the four samples with degradation or low tumor cell content issues) and all other samples in the cohort using the independent t-test function *ttest_ind()* of scipy v1.11.4. Benjamini-Hochberg multiple testing correction was performed with the *fdrcorrection()* method of statsmodels v0.14.2.

#### ATR substrate correlation analysis

For each phosphopeptide detected in at least 1 QC channel for 67% of batches (i.e. those for which bridge normalization was performed) and quantified in >200 patients, the Pearson correlation was calculated between the ATR TOPAS substrate phosphorylation subscore and the protein expression corrected abundances (as described in the TOPAS score section above) of the phosphopeptide across patients. Known ATR/ATM/PRKDC substrates were extracted from PhosphoSitePlus and *Bayer et al.*^15^ and phosphopeptides with the pSQ/pTQ motif were marked. Phosphopeptides that had a Pearson correlation >0.5, had the pSQ/pTQ motif and were not annotated as an ATR/ATM/PRKDC substrate in PhosphoSitePlus were considered as potential new substrates of these kinases.

