## Extended Data Figures for "TOPAS: phosphoproteome data analysis and decision support platform for molecular tumor boards"

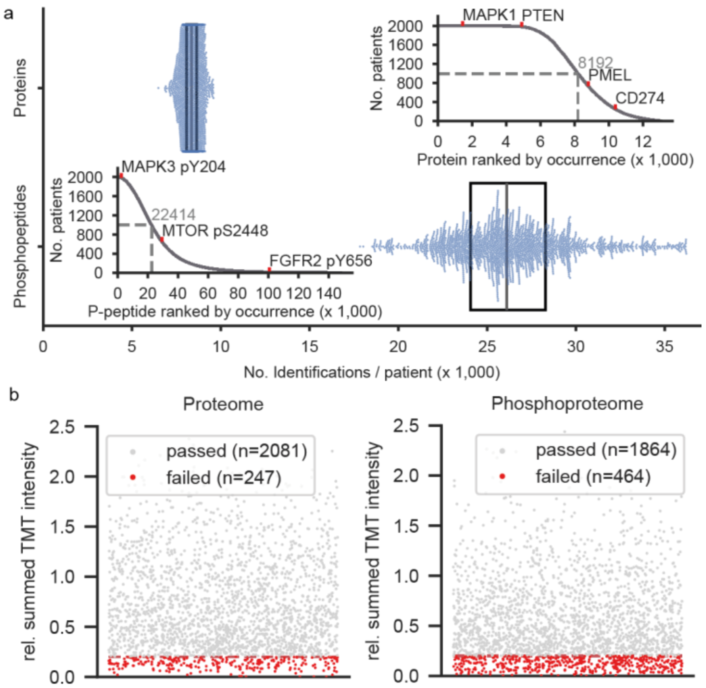


**Extended Data Fig. 1: Phosphoproteome coverage and quality control.** (a) Swarm plots with boxplots overlaid showing the number (and median) of identified proteins and phosphopeptides in the 1,998 patient samples. The insets show a ranked list of proteins or phosphopeptides, respectively, sorted by how many patient samples they were detected in (occurrence). (b) Strip plots of summed TMT reporter intensity for each patient sample relative to the mean of the QC channels in the same TMT batch. A ratio of 5-fold (0.2) below the mean of the QC channels was deemed of insufficient quality and removed from the cohort.

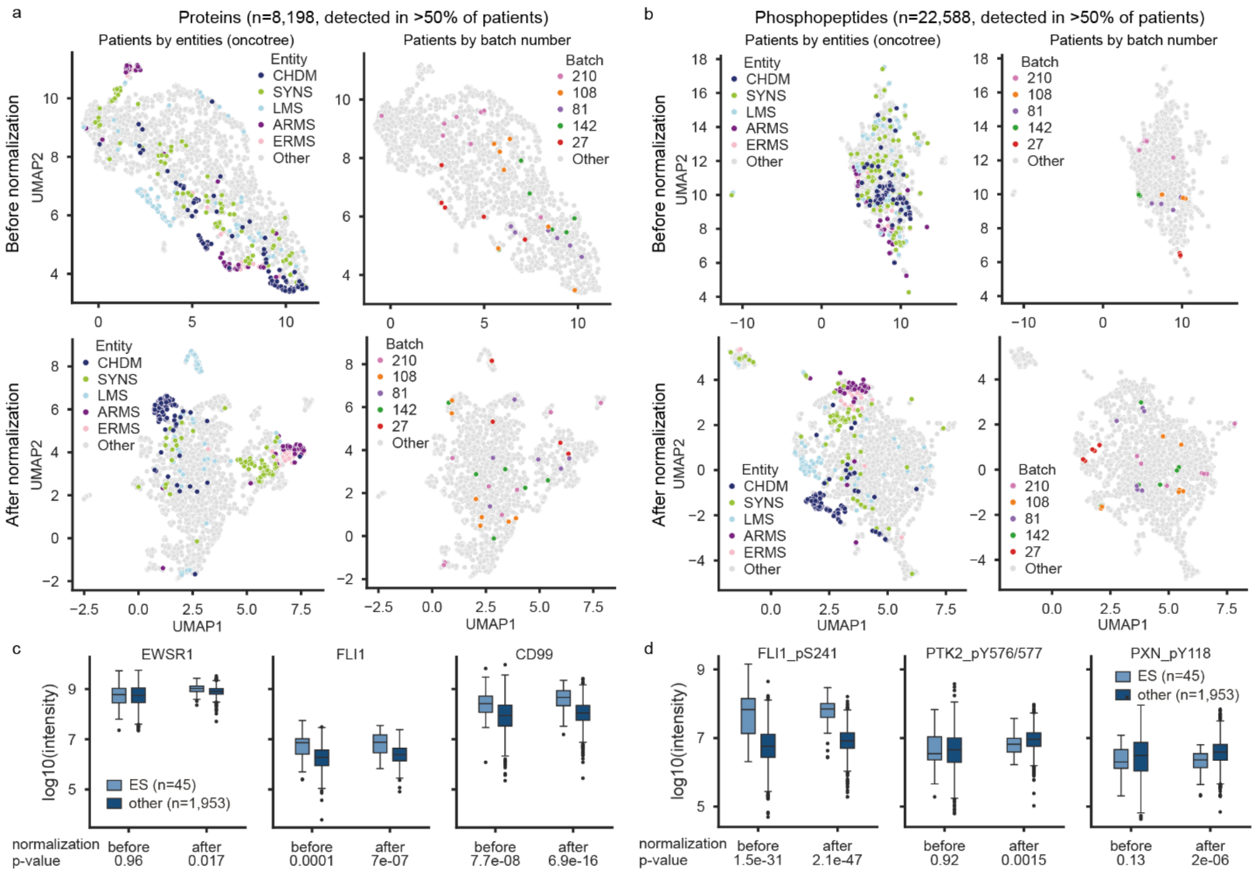


**Extended Data Fig. 2: Phosphoproteome data normalization.** (a) UMAP analysis of patient proteomes colored by entity (left panel) or batch number (right panel) before (top row) and after (bottom row) normalization. (b) same as (a) but for patient phosphopeptides (c) Box plots of protein abundance of three known biomarkers for Ewing Sarcoma (ES) and detected in ES patients in this study: EWSR1 (n=45), FLI1 (n=23), CD99 (n=45) vs all other patients in the cohort (EWSR1: n=1,953, FLI1: n=455, CD99: n=1,952) before and after normalization. (d) Box plots of phosphopeptide abundance of three putative biomarkers for ES: FLI1 pS241 (n=45), PTK2 pS576/577 (n=44), PXN pY118 (n=43) vs all other patients in the cohort (FLI1 pS241: n=1,864, PTK2 pS576/577: n=1,930, PXN pY118: n=1,842) before and after normalization. ARMS, Alveolar Rhabdomyosarcoma, ERMS, Embryonal Rhabdomyosarcoma, CHDM, Chordoma, LMS, Leiomyosarcoma, SYNS, synovial sarcoma

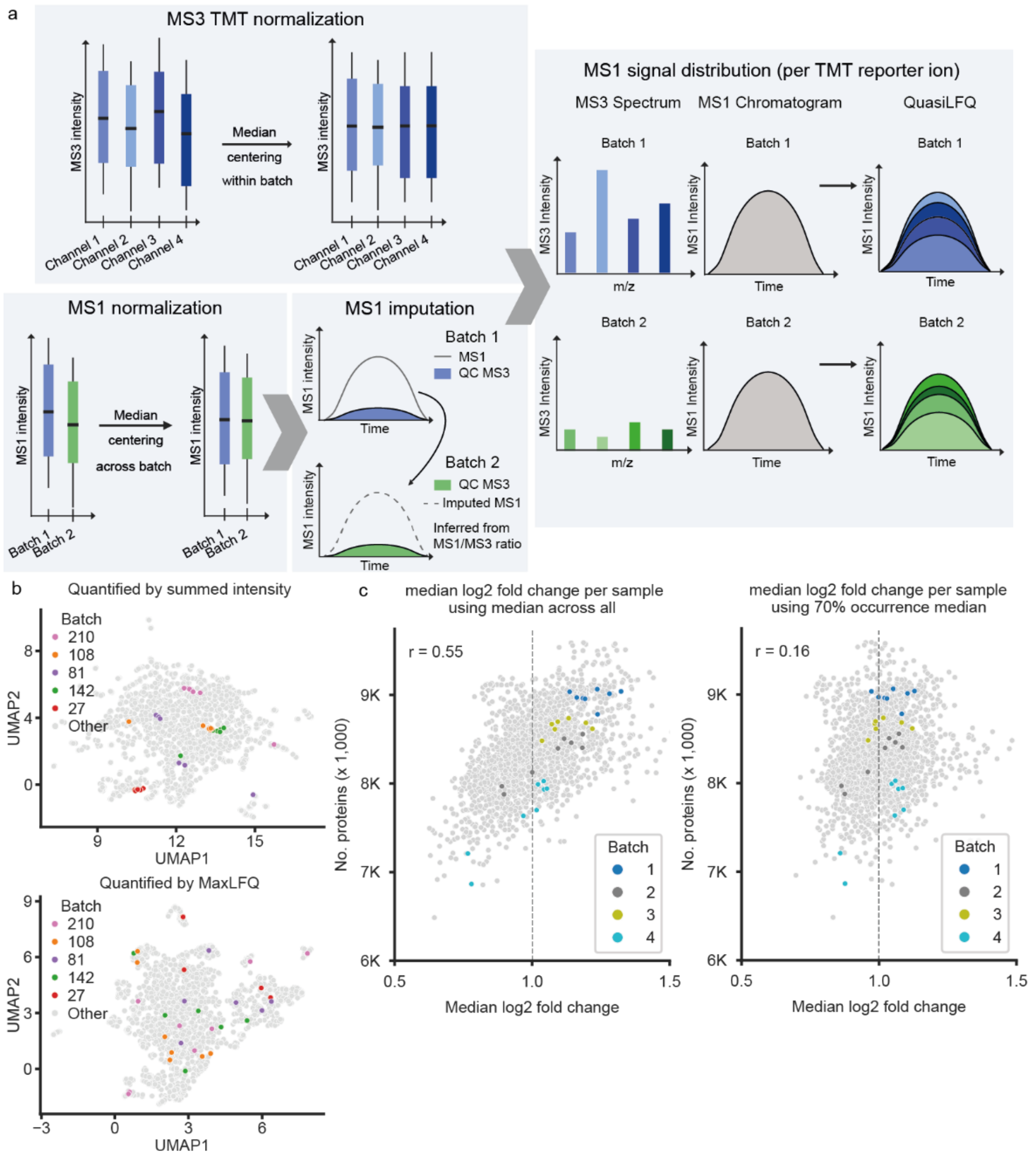


**Extended Data Fig. 3: Reducing batch effects using simple methods.** (a) Graphical depiction of the data normalization process. Median centering of TMT reporter ion intensities recorded in the MS3 spectrum was used to normalize within each TMT batch (top left panel). Median centering of peptide intensities recorded in MS1 spectra was used for normalization between batches. This required imputation of a small fraction of MS1 peptide intensities using the expected ratio of MS1 vs MS3 signal intensities of the QC samples present in every batch (bottom left panel). This was followed by the computation of quasi-LFQ intensities by distributing the MS1 intensity (the composite signal of all patients and QC samples in one TMT batch) according to the relative contribution of the respective TMT MS3 intensities that each represent a single patient or QC sample (right panel). (b) UMAP analysis of patient proteomes comparing the summed peptide intensity approach for protein quantification (top panel) to the MaxLFQ algorithm. (c) Scatter plots of median of log2 fold changes of all protein abundances per patient (x-axis) vs the number of identified proteins (y-axis) in that patient. For the left plot, the median peptide intensity used for median centering normalization of a batch (MS1, see panel a) was calculated using all peptides identified in this batch. For the right plot, only peptides found in at least 70% of all batches were used.

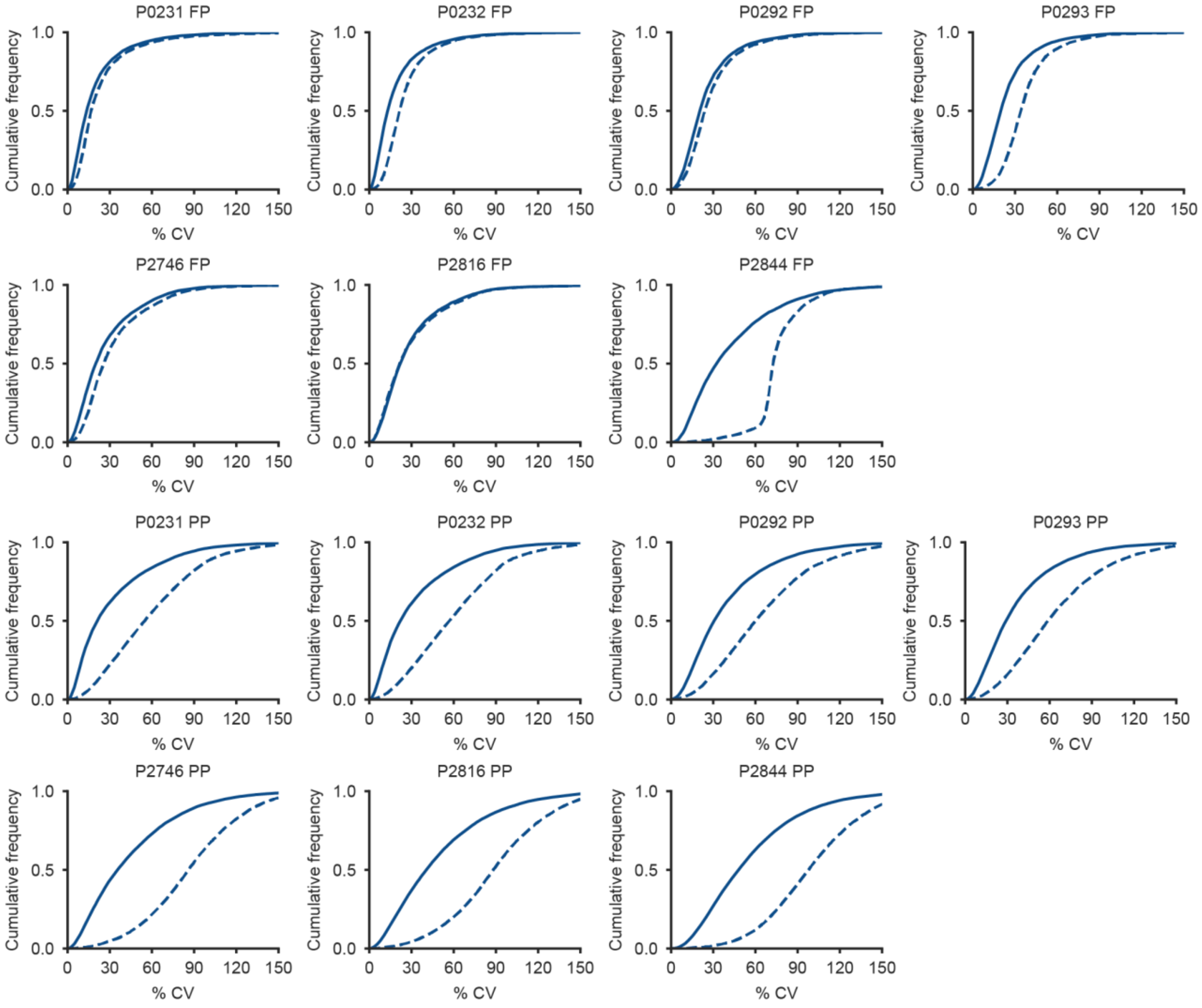


**Extended Data Fig. 4**: **Data normalization across sample batches reduces variation in phosphoproteomes.** Cumulative frequency plots showing the effect of data normalization (dashed line before and solid line after normalization) on quantitative precision (coefficient of variation, CV) of full workflow replicates of 7 patients (n=3 for each patient). FP: full proteome, PP: phosphoproteome.

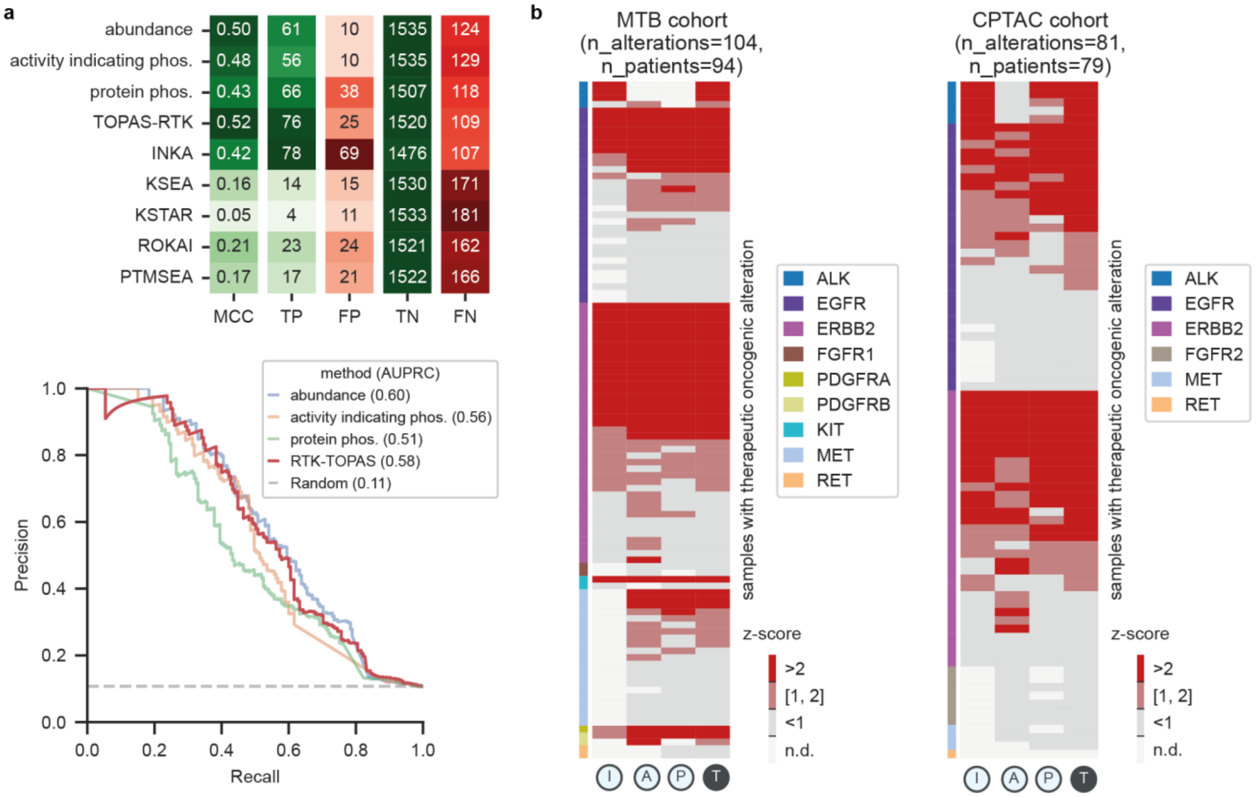


**Extended Data Fig. 5**: **Comparison of kinase activity inference methods.** (a) Heatmap of classification evaluation metrics for a range of kinase activity inference methods (top). Precision-recall curve comparing RTK-TOPAS to its constituent scores (bottom). (b) Heatmap of RTK-TOPAS scores and their constituent sub-scores (I: activity indicating phosphorylation, A: abundance, P: protein phosphorylation, T: TOPAS-RTK) for patients carrying genome alterations in certain RTKs (colored bars) from the MTB (left panel) and CPTAC (right panel) cohorts. Each row is a patient with a therapeutically relevant oncogenic alteration. MCC=Matthew's correlation coefficient, TP=True positive, FP=False positive, TN=True negative, FN=False negative, AUPRC=Area Under the Precision-Recall Curve.

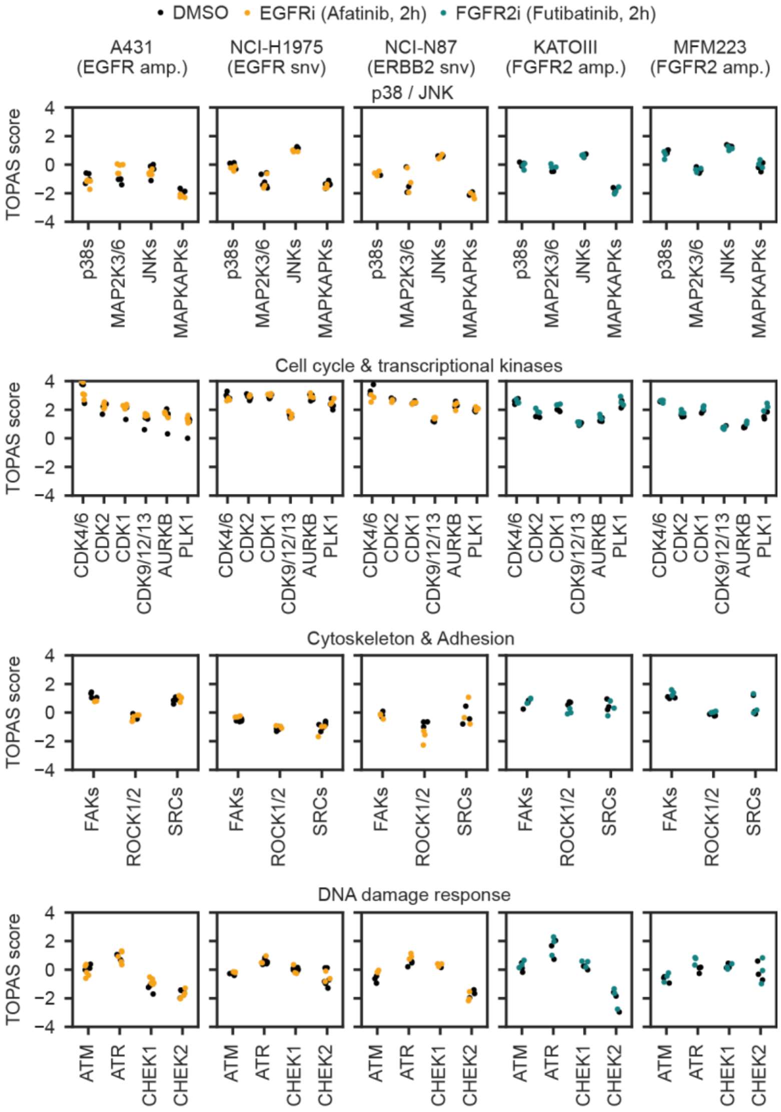


**Extended Data Fig. 6**: **ICK-TOPAS scores in response to drug perturbation.** Scatter plots of ICK-TOPAS scores for five cell lines with known RTK oncogenic driver signals in which four indicated pathways (p38/JNK, cell cycle and transcription, cytoskeleton and adhesion, DNA damage response) were unaffected by drug perturbation.

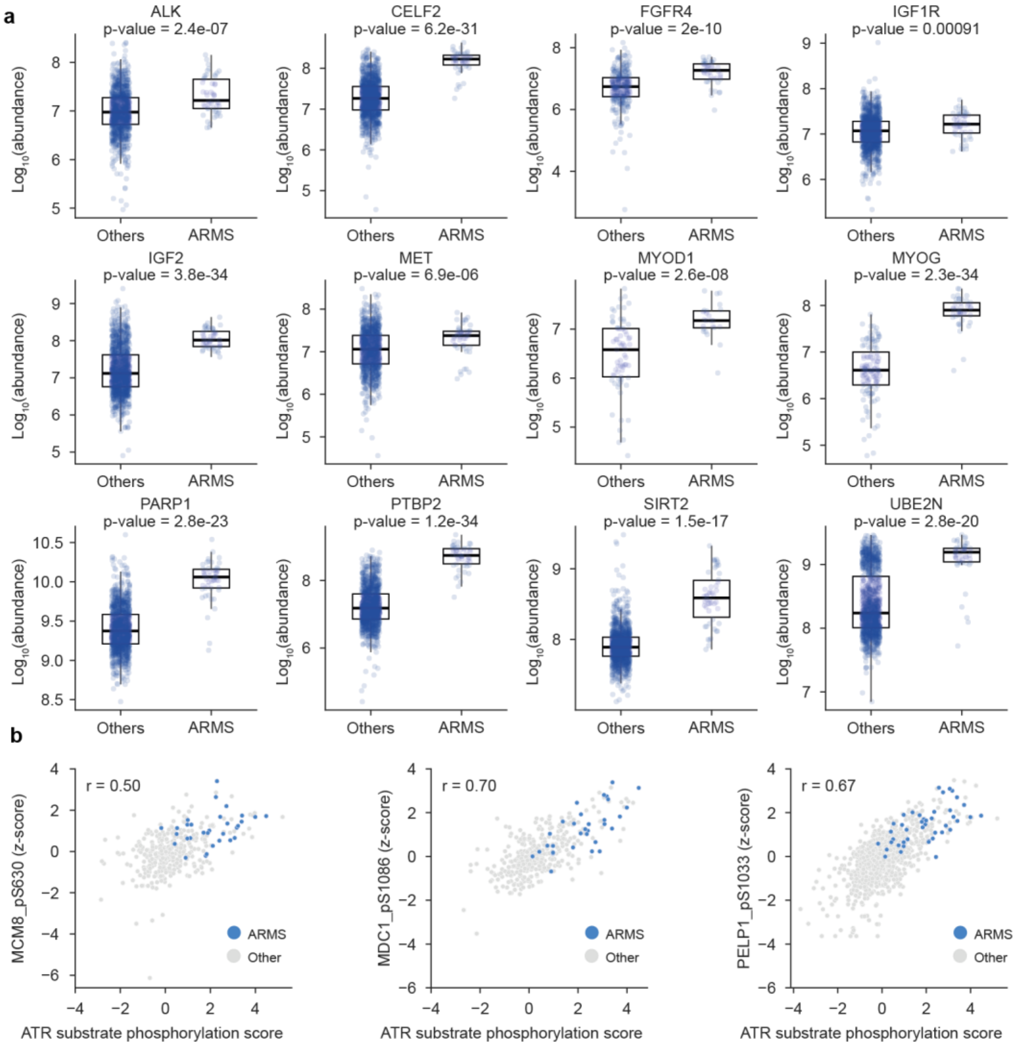


**Extended Data Fig. 7**: **Differential protein abundance and elevated ATR activity in ARMS patients.** (a) Swarm plots with overlaid box plots of exemplary proteins that are differentially abundant between ARMS patients (n=42) and the rest of the cohort (n=1,956). (b) Examples for abundance correlation plots of annotated (PELP1_pS1033) and previously unannotated (FUBP_pS569, POP1_pS367) phospho-sites (y-axis) carrying a SQ/TQ ATR substrate motif vs ATR substrate phosphorylation score (x-axis) computed for patient samples. ARMS samples are highlighted in blue, all other samples in grey.
